# Chromatin crosstalk between HDA19 and NuA4 sets thresholds for stress gene activation in *Arabidopsis*

**DOI:** 10.64898/2026.01.12.699102

**Authors:** Wojciech Dziegielewski, Anna Bieluszewska, Tomasz Bieluszewski, Michal Krzyszton, Alexandre Pelé, Maja Szymanska-Lejman, Anna Wilhelm, Szymon Swiezewski, Piotr A. Ziolkowski

## Abstract

Histone acetylation by the plant NuA4 complex promotes high expression of growth-related genes while preventing gene body H2A.Z depletion and spurious activation of stress-responsive loci. How this NuA4-dependent chromatin state is modulated by histone deacetylases (HDACs) in plants remains unclear. Here, we investigate the interplay between NuA4 and HDACs in *Arabidopsis* by generating a collection of HDAC loss-of-function mutants in NuA4-proficient and NuA4-deficient backgrounds. Loss of individual HDACs did not rescue the severe growth defects of the NuA4(−) mutant; instead, most combinations further aggravated the phenotype, indicating that HDACs predominantly support rather than counteract NuA4 function. Focusing on the *hda19-5^Atepl1b^*allele, we show that loss of HDA19 causes strong developmental defects and constitutive activation of biotic stress–responsive genes, but not canonical heat-response genes. 3′ RNA-seq and ChIP-seq reveal that upregulated genes in *hda19-5^Atepl1b^*display increased H3K9ac and H2A.Zac with largely unchanged gene-body H2A.Z, and substantially overlap with genes induced in the NuA4-null *Atepl1-2* mutant and in wild-type plants exposed to elevated temperature. Integration of our datasets with published HDA19 ChIP-seq maps shows that H2A.Zac is selectively increased at HDA19-bound H2A.Z peaks in *hda19-5^Atepl1b^*, consistent with a direct role for HDA19 in H2A.Z deacetylation. However, elevated H2A.Zac is neither necessary nor sufficient for transcriptional activation, whereas changes in H3K9ac correlate strongly with gene induction. We propose that NuA4 and HDA19 cooperate to tune chromatin at stress-related loci, with H3K9 acetylation as the primary driver of transcription and H2A.Z acetylation acting as a modulatory mark that shapes stress gene responsiveness.

## Introduction

Plants are continuously exposed to adverse environmental conditions but cannot escape unfavourable weather or pathogen attack (Doerner, 2020). Instead, they rely on elaborate sensing, signalling, and defence mechanisms to cope with biotic and abiotic stresses (War et al., 2012; Pierik and Testerink, 2014). Since stress responses compete for the resources with growth, their dynamic coordination is essential. Our previous work has implicated the histone variant H2A.Z and the NuA4 histone acetyltransferase complex as key regulators of this balance (Sura et al., 2017; Bieluszewski et al., 2022a).

H2A.Z is typically enriched at the +1 nucleosome downstream of the transcription start site (TSS), but can also be broadly distributed across gene bodies (Zilberman et al., 2008). Depending on its genomic localization and posttranslational modification state, H2A.Z can support either gene activation or repression (Coleman-Derr and Zilberman, 2012; Sura et al., 2017). High H2A.Z occupancy at the +1 nucleosome is associated with active genes and is thought to lower the energetic barrier for RNA Polymerase II (RNAPII) during transcription initiation (Weber et al., 2014). NuA4-mediated acetylation of H2A.Z at this position promotes transcription of photosynthesis-related genes (Bieluszewski et al., 2022a). By contrast, many environmentally responsive genes show elevated levels of gene body H2A.Z, which is proposed to keep them repressed under non-inducing conditions (Coleman-Derr and Zilberman, 2012; Sura et al., 2017). Consistent with this model, active eviction of H2A.Z accompanies induction of several stress-regulated genes (Zander et al., 2019; Willige et al., 2021; Xue et al., 2021). More recent data indicate that the relationship between gene body H2A.Z and transcriptional repression is more complex and depends on specific posttranslational modifications of H2A.Z and other histones (Gómez-Zambrano et al., 2019; Sijacic et al., 2025; Zhu et al., 2025).

We previously showed that NuA4 not only acetylates H2A.Z but also promotes its nucleosomal deposition in *Arabidopsis*, in agreement with earlier findings in yeast (Altaf et al., 2010; Cheng et al., 2015; Bieluszewski et al., 2022a). In NuA4-null mutants, stress-responsive genes display reduced H2A.Z occupancy and increased transcript levels (Bieluszewski et al., 2022a). These observations suggest that, under non-inducing conditions, NuA4 activity that supports H2A.Z deposition and transcriptional competence must be continuously counteracted by histone deacetylation over the bodies of stress-responsive genes, thereby maintaining H2A.Z in a repressive mode.

*Arabidopsis* encodes 18 histone deacetylases (HDACs) classified into three subfamilies (Alinsug et al., 2009). Twelve of them are Zn²⁺-dependent RPD3/HDA1-type enzymes, which are further subdivided into Classes I, II, and IV. The remaining HDACs belong either to the plant-specific HD-Tuin family (four members) or to the Sirtuin (SIR) family (two members) (Ma et al., 2013). This expanded HDAC repertoire suggests not only functional redundancy but also specialization and potentially antagonistic roles in stress regulation.

Indeed, multiple studies have demonstrated that HDACs affect diverse biological processes in plants, including stress responses (Luo et al., 2012; Morończyk et al., 2022; Wu et al., 2008; Liu et al., 2017; Kim et al., 2013; Zheng et al., 2020). For example, Class I mutants such as *hda19* and *hda9* display increased salt tolerance, whereas the *hda5/14/15/18* quadruple mutant is hypersensitive to salt stress (Ueda et al., 2017; Zheng et al., 2020). Similarly, *hda6* mutants are more tolerant to drought, whereas *hda15* mutants are hypersensitive (Kim et al., 2017; Lee and Seo, 2019). In temperature responses, *hda9* and *hda19* mutants show warm-insensitive phenotypes, in contrast to *hda15* (Shen et al., 2019). Plant-specific HD-Tuins also contribute to drought and heat stress responses (Han et al., 2016; Buszewicz et al., 2016). Consistently, the expression of genes involved in heat, drought, and osmotic stress is altered across several *hdac* mutant backgrounds (van der Woude et al., 2019; Shen et al., 2019; Han et al., 2016).

To identify which *Arabidopsis* HDACs antagonize NuA4-dependent acetylation, we generated individual *hdac* knockouts using CRISPR/Cas9 in a NuA4-deficient (*Atepl1a-1 -/- Atepl1b-1 -/-*, hereafter *Atepl1-1*) background. We hypothesized that loss of a direct NuA4 antagonist would restore the activity of growth-related genes, as reported in yeast (Lin et al., 2008). Contrary to this expectation, most triple *Atepl1-1 hdac* homozygous mutants showed an aggravated *Atepl1-1* phenotype. We therefore expanded our analysis to double mutants (*Atepl1a-1 +/+ Atepl1b-1 -/- hdac -/-*), which approximate HDAC loss in an otherwise wild-type background due to the functional redundancy between EPL1A and EPL1B. Among these, *Atepl1a-1 +/+ Atepl1b-1 -/-hda19-5 -/-* (*hda19-5^Atepl1b^*) exhibited the strongest reduction in rosette size. Transcriptome profiling of *hda19-5^Atepl1b^*revealed upregulation of stress-responsive genes that are normally induced at elevated ambient temperature. ChIP-seq analysis further showed increased H3K9ac and H2A.Zac levels at a subset of these loci. Together, our data support a model in which HDA19 removes acetyl groups from H3K9 and H2A.Z at stress-related genes, thereby preventing their inappropriate activation under physiological growth conditions.

## Results

### Individual histone deacetylase knockouts aggravate NuA4(−) phenotype

To test whether disruption of individual histone deacetylases can restore the balance between histone acetylation and deacetylation and thereby rescue a mutant lacking a functional NuA4 complex, we generated a uniform collection of novel CRISPR/Cas9 *hdac* alleles in the *Atepl1-1* background. *Atepl1-1* is a loss-of-function NuA4 mutant, corresponding to a double mutant lacking the paralogous NuA4 subunits AtEPL1A and AtEPL1B. Using a high-efficiency CRISPR/Cas9 protocol (Bieluszewski et al., 2022b), we obtained loss-of-function deletions in 15 HDAC genes, excluding *HDA8* and the pseudogenes *HDA10* and *HDA17* (de Rooij et al., 2021). Most mutations targeted the first exon of the respective *HDAC* gene, resulting in frameshifts and premature stop codons (Supplementary Fig. 1-13; Supplementary Table 1).

Because *Atepl1-1* double homozygotes are completely sterile, the novel histone deacetylase alleles (hereafter collectively referred to as *hdac* alleles) were first generated in the sesquimutant *Atepl1a-1 +/− Atepl1b-1 -/-* background. After sequence verification of heritable loss-of-function alleles, we isolated combinations of *Atepl1-1* double homozygotes with homozygous mutations in 13 histone deacetylases: *HDT1–HDT4, HDA2, SRT1, SRT2, HDA6, HDA7, HDA9, HDA14, HDA15, HDA19* and a quadruple mutation affecting the tandemly arranged *HDA5* and *HDA18* (*Atepl1a-1 Atepl1b-1 hda5 hda18*) (Supplementary Fig. 14). Although we successfully generated a homozygous *hda6* null allele in the *Atepl1-1* sesquimutant background, no corresponding triple homozygous mutants could be recovered, indicating that *hda6* is lethal in the absence of EPL1 (χ² test, *p* = 3.26 × 10⁻¹⁷; n = 102).

The *Atepl1-1* double mutant displays a strongly reduced rosette diameter, which can be explained by decreased transcription of growth-related genes due to reduced histone acetylation, together with increased activity of stress-related genes associated with decreased H2A.Z occupancy (Bieluszewski et al., 2022a). We initially hypothesized that loss of an HDAC antagonistic to NuA4 would compensate for the absence of NuA4 and rescue the *Atepl1-1* phenotype. Instead, several *hdac* mutations aggravated the *Atepl1-1* phenotype, and none alleviated it (Supplementary Fig. 14; Supplementary Table 2). To account for potential functional redundancy between histone deacetylases from different classes (Shen et al., 2019), we also generated a quadruple *Atepl1-1 hda15 hda19* mutant. However, its phenotype was similar to that of the triple *Atepl1-1 hda19* mutant (Supplementary Fig. 14; Supplementary Table 2). These observations suggest that transcriptional upregulation of stress-responsive genes in the triple mutants outweighs any positive effect of increased histone acetylation on growth-related genes that are hypoacetylated in the *Atepl1-1* background.

Because *Atepl1b-1* plants do not exhibit visible vegetative or generative defects due to functional redundancy between EPL1A and EPL1B (Supplementary Fig. 15; Supplementary Table 3), we used double homozygous *Atepl1b-1 -/- hdac -/-* mutants to assess the impact of *hdac* mutations on growth in an otherwise wild-type–like background (Fig. 1A; Supplementary Table 3). Among the mutants analysed, *hda19-5 Atepl1b* (*hda19-5^Atepl1b^*) showed the most pronounced size reduction, with more than a twofold decrease in rosette diameter compared to wild-type plants (mean = 2.75 cm; Welch’s t-test, *p* = 3.82 × 10⁻⁷). The *hda19-5^Atepl1b^*mutant exhibited narrow cotyledons and reduced fertility, consistent with previous descriptions of *hda19* mutants (Fig. 1B–C) (Ning et al., 2019; Tanaka et al., 2007). The generated *HDA19* allele carries a 1 bp deletion followed by a 47 bp deletion in the first exon, leading to a frameshift mutation (Fig. 1D).

**Fig. 1.**
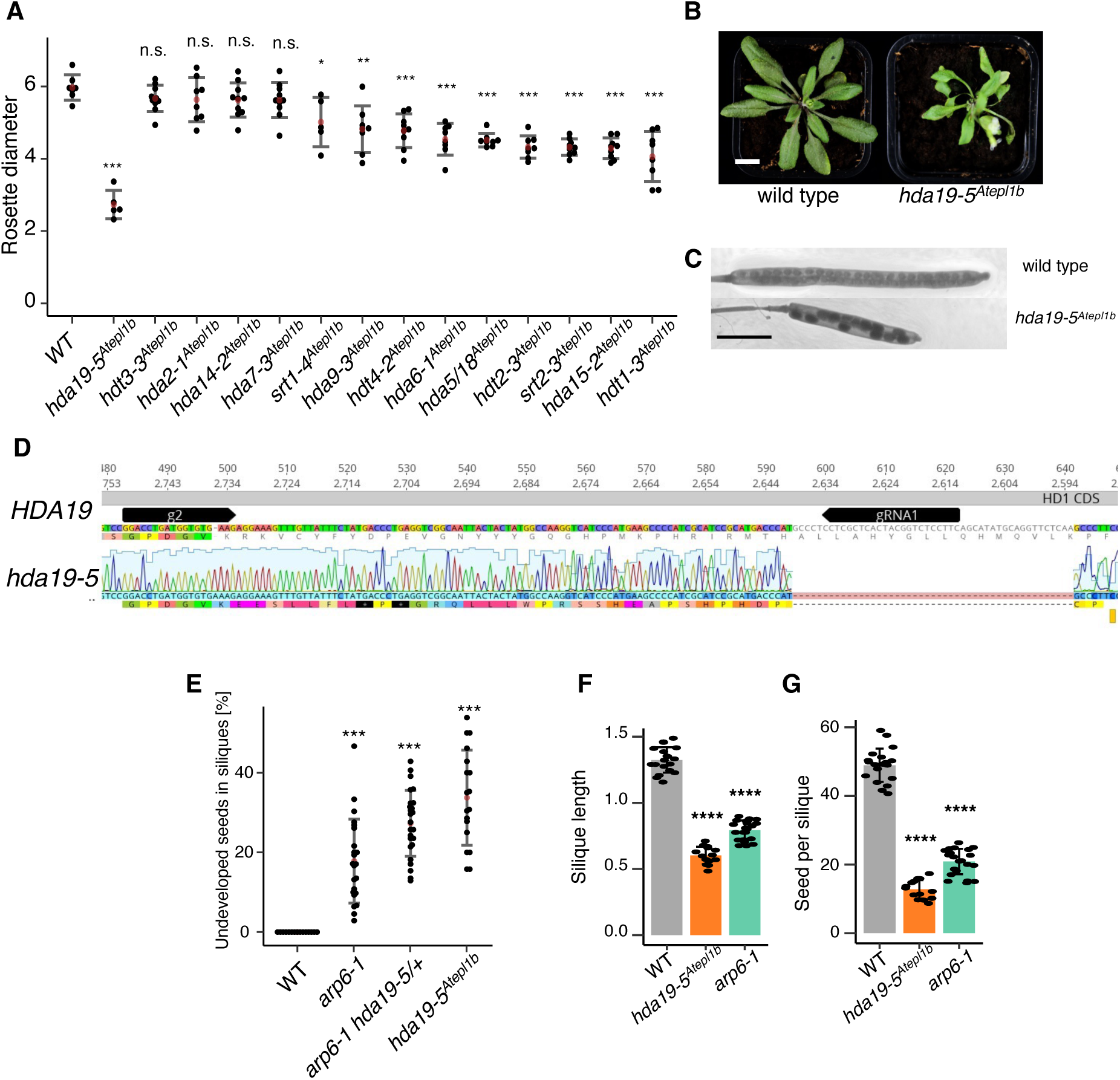
Mutants of *hda19-5^Atepl1b^* display developmental and fertility defects. **(A)** Comparison of rosette size in *hdac****^Atepl1b^*** mutants. Bars represent mean ± SD. Statistical significance was assessed using a two-sided Welch’s t-test. Significance levels: *p* < 0.05 *, *p* < 0.01 **, *p* < 0.001 ***, *p* < 0.0001 ****, non-significant (n.s.). **(B)** Representative images of the *hda19-5****^Atepl1b^*** mutant and wild-type (WT) plants. Scale bar = 1 cm. **(C)** Representative siliques of WT and *hda19-5****^Atepl1b^*** plants. Scale bar = 20 mm. **(D)** Sequence of the *hda19-5* allele showing a 1 bp deletion followed by a 47 bp deletion in the first exon, resulting in a frameshift. **(E)** Fertility analysis of *hda19-5 +/−* × *arp6-1 -/-*, *hda19-5****^Atepl1b^***, and *arp6-1* plants. Statistical significance was determined using a two-sided Welch’s t-test, with significance levels indicated as in (A). **(F, G)** Quantification of silique length (F) and seed set (G) in *hda19-5****^Atepl1b^*** and *arp6-1* mutants compared with WT. Statistical significance was determined using a two-sided Welch’s t-test, with significance levels indicated as in (A).

In summary, our genetic analyses suggest that histone deacetylases predominantly support NuA4’s indirect role in transcriptional repression of stress responses, rather than simply antagonizing its function in transcriptional activation of growth-related genes.

### Mutants of *HDA19* and *ARP6* exhibit similar phenotypes and synthetic lethality

Plants lacking the ARP6 subunit of the H2A.Z-depositing SWR1 complex display elevated stress responses without the severe downregulation of the photosynthetic apparatus observed in NuA4(−) mutants (Sura et al., 2017). We previously showed that loss of ARP6 is synthetically lethal with *Atepl1-1* (Bieluszewski et al., 2022a). To assess the impact of *hdac* mutations in the absence of nucleosomal H2A.Z, we tried to generate the *arp6-1 hda19-5* double mutant (with functional NuA4). Among 100 progeny from the *hda19-5* +/− *arp6-1* -/- F_2_ plants, no double homozygous plants were recovered, indicating synthetic lethality (χ² test, *p* = 1.36 × 10⁻⁸; Supplementary Table 4). The siliques of *hda19-5* +/− *arp6-1* -/- plants contained fewer developed seeds than *arp6-1* -/- alone, but more than *hda19-5^Atepl1b^*, suggesting distinct yet indispensable roles of HDA19 and ARP6 in plant reproduction (Fig. 1E; Supplementary Table 5).

To better understand the basis of this synthetic lethality, we compared the phenotypes of the mutants. Both *arp6-1* and *hda19-5^Atepl1b^*exhibited reduced silique length, decreased seed set, and smaller rosettes relative to wild-type (WT) plants (Fig. 1F–G; Supplementary Table 5). In addition, *arp6-1* showed significantly reduced pollen density compared with WT and *hda19-5^Atepl1b^*, consistent with previous reports (*p* = 7.47 × 10⁻⁵; Supplementary Fig. 16; Supplementary Table 6) (Deal et al., 2005).

Cytogenetic analysis revealed that male meiosis proceeds normally in both *arp6-1* and *hda19-5^Atepl1b^*. Examination of 180 pollen mother cells per genotype showed no chromosome bridges or fragmentation (Supplementary Fig. 17; Supplementary Table 7). These observations suggest that fertility defects in the mutants likely arise from post-meiotic processes or female-specific defects, consistent with recent findings for *hda19* mutants (Manrique et al., 2024).

Given that many stress-related genes are upregulated in *arp6-1* due to global H2A.Z loss (Dai et al., 2017; Potok et al., 2019; Sura et al., 2017), we hypothesized that H2A.Z dynamics might also be perturbed in the *hda19-5^Atepl1b^*background. Such disruption could account for the phenotypic similarities between these mutants and may underlie their synthetic lethality.

### Transcriptomic profile of *hda19^Atepl1b^* mimics the effects of physiological and genetic depletion of H2A.Z

To understand the basis of the phenotypic similarity and synthetic lethality between *hda19* and *arp6* mutants, we quantitatively analysed gene expression changes in *hda19-5^Atepl1b^*using 3′ RNA-seq, which selectively sequences the 3′ ends of cDNA libraries (Krzyszton et al., 2024; Alpern et al., 2019). Because previous studies showed that H2A.Z is removed from nucleosomes genome-wide in response to elevated temperature, we also included wild-type and *hda19-5^Atepl1b^* plants grown at 27°C to compare the effects of physiological and genetic H2A.Z depletion (Shen et al., 2019; Ning et al., 2019).

In *hda19-5^Atepl1b^* grown at 22°C, we identified 656 upregulated (log₂FC > 1) and 405 downregulated genes (log₂FC < −1) relative to wild type at 22°C (Fig. 2A; Supplementary Table 8). Wild-type plants grown at 27°C showed 962 upregulated and 929 downregulated genes compared with wild type at 22°C. Although principal component analysis (PCA) did not cluster *hda19-5^Atepl1b^* at 22°C together with wild type at 27°C (Supplementary Fig. 18A), previous reports on the role of HDA19 in temperature responses (Shen et al., 2019) led us to hypothesize that similar gene sets might be activated in these conditions. Indeed, 66% (437) of the genes upregulated in *hda19-5^Atepl1b^* at 22°C were also upregulated in wild type at 27°C (hypergeometric test, *p* < 2 × 10⁻¹⁶; Fig. 2B). These shared genes displayed comparable fold changes in both contexts (Fig. 2C) and were enriched for Gene Ontology (GO) terms related to responses to biotic stimuli (Fig. 2D; Supplementary Table 9). Notably, despite the extensive overlap, genes associated with heat response were specifically induced in wild type at 27°C but not in *hda19-5^Atepl1b^* at 22°C (Supplementary Table 9), suggesting that HDA19 predominantly represses stress genes linked to pathogen responses rather than those involved in abiotic heat stress.

**Fig. 2.**
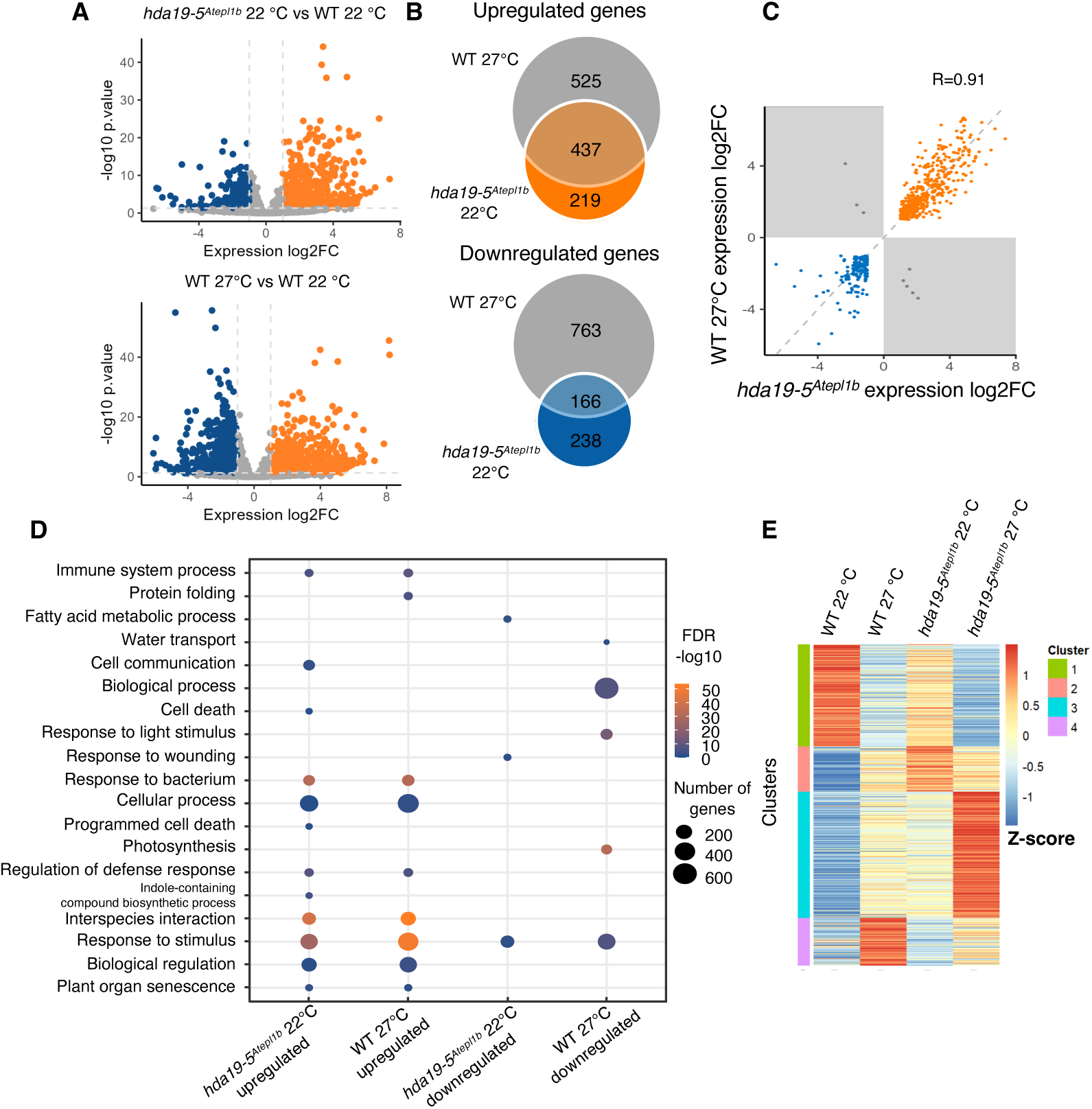
3′ RNA-seq analysis of *hda19-5^Atepl1b^* and wild-type plants grown at elevated temperature. **(A)** Volcano plots showing differentially expressed genes in *hda19-5****^Atepl1b^*** at 22°C (top) and wild type (WT) grown at 27°C (bottom), each compared with WT at 22°C. The x-axis shows log₂ fold change, and the y-axis shows statistical significance (−log₁₀ *p*-value). Up-regulated genes (log₂FC > 1) are shown in orange, downregulated genes (log₂FC < −1) in blue. **(B)** Venn diagrams showing overlaps between upregulated (top) and downregulated (bottom) genes in *hda19-5****^Atepl1b^*** at 22°C, *hda19-5****^Atepl1b^*** at 27°C, and WT at 27°C. Significance of the overlaps was assessed using a hypergeometric test. **(C)** Comparison of fold changes for genes commonly up- or downregulated in *hda19-5****^Atepl1b^*** at 22°C and WT at 27°C. The correlation coefficient was calculated using Pearson’s correlation (p < 2.2 × 10⁻¹⁶). **(D)** Gene Ontology (GO) enrichment analysis for genes dysregulated in *hda19-5****^Atepl1b^*** at 22°C and WT at 27°C. The x-axis indicates the experimental condition, and the y-axis shows significantly enriched GO terms. Redundant GO terms were reduced using REVIGO to retain representative categories. Dot size indicates the number of genes per category, and colour encodes statistical significance. **(E)** Heatmap of Z-scored expression values for the 1000 most variable genes across WT and *hda19-5****^Atepl1b^*** plants grown at 22°C and 27°C, clustered by *k*-means.

Downregulated genes showed a similar, albeit weaker, pattern of overlap: 41% (166) of the genes downregulated in *hda19-5^Atepl1b^* at 22°C were also downregulated in wild type at 27°C (Fig. 2B–C). These genes were associated with multiple responsive and metabolic GO categories (Fig. 2D; Supplementary Table 9).

When we compared *hda19-5^Atepl1b^* and wild type directly at 27°C, 594 genes were upregulated and 341 were downregulated in *hda19-5^Atepl1b^*(Supplementary Fig. 18B). However, direct differential analysis between *hda19-5^Atepl1b^*at 22°C and 27°C is confounded by the fact that normalization is performed against the corresponding wild-type datasets at each temperature. As a result, genes already misregulated in *hda19-5^Atepl1b^* at 22°C may not appear as differentially expressed between temperatures, even if their absolute expression levels change further at 27°C.

To circumvent this limitation, we focused on the 1000 most variable genes across all samples (wild type and *hda19-5^Atepl1b^* at 22°C and 27°C) and performed *k*-means clustering of their expression profiles. This analysis resolved four distinct expression clusters (Fig. 2E). Genes in cluster 1, enriched for photosynthesis- and growth-related functions, were highly expressed in both wild type and *hda19-5^Atepl1b^* at 22°C (Supplementary Fig. 18C–D). Cluster 2 comprised stress-response genes that were most strongly induced in *hda19-5^Atepl1b^* at 22°C, indicating constitutive activation of stress pathways in the mutant under physiological conditions. Cluster 3 contained responsive genes that were already moderately upregulated (i.e., “primed”) in *hda19-5^Atepl1b^* at 22°C, but reached maximal expression in *hda19-5^Atepl1b^* at 27°C, suggesting additive effects of environmental cues and loss of HDA19 activity. Finally, the relatively small cluster 4 included genes that were most strongly induced specifically in wild-type plants at 27°C (Supplementary Fig. 18C–D), representing a subset of temperature-responsive genes whose activation does not depend on HDA19. Dysregulation of selected genes identified by 3′ RNA-seq was validated by RT-qPCR (Supplementary Fig. 19).

### Stress-response genes display H2A.Z and H3K9 hyperacetylation in *hda19-5^Atepl1b^*

Previous studies have primarily implicated HDA19 in the deacetylation of histone H3 and H4 (Zhou et al., 2013; Manrique et al., 2024; Gao et al., 2015; Liu et al., 2025). However, we hypothesized that HDA19 may also target the H2A.Z variant, thereby contributing to the repression of stress-response genes (Temman et al., 2023; Ueda et al., 2017; Zhou et al., 2013). To test this, we performed ChIP-seq for H3K9ac, H2A.Zac, and total H2A.Z in the *hda19-5^Atepl1b^* background.

We first examined global levels of these marks across genes expressed in wild-type plants grown at 22°C (n = 19,256; genes with >10 detected transcripts in the 3′ RNA-seq dataset). Metagene profiles revealed a slight but significant decrease in H3K9ac at the +1 nucleosome in *hda19-5^Atepl1b^*compared with wild type (two-sided unpaired Wilcoxon test, *p* = 0.048; Fig. 3A). In contrast, average H2A.Zac and total H2A.Z levels did not differ markedly between *hda19-5^Atepl1b^* and wild type for the majority of genes (Fig. 3A).

**Fig. 3.**
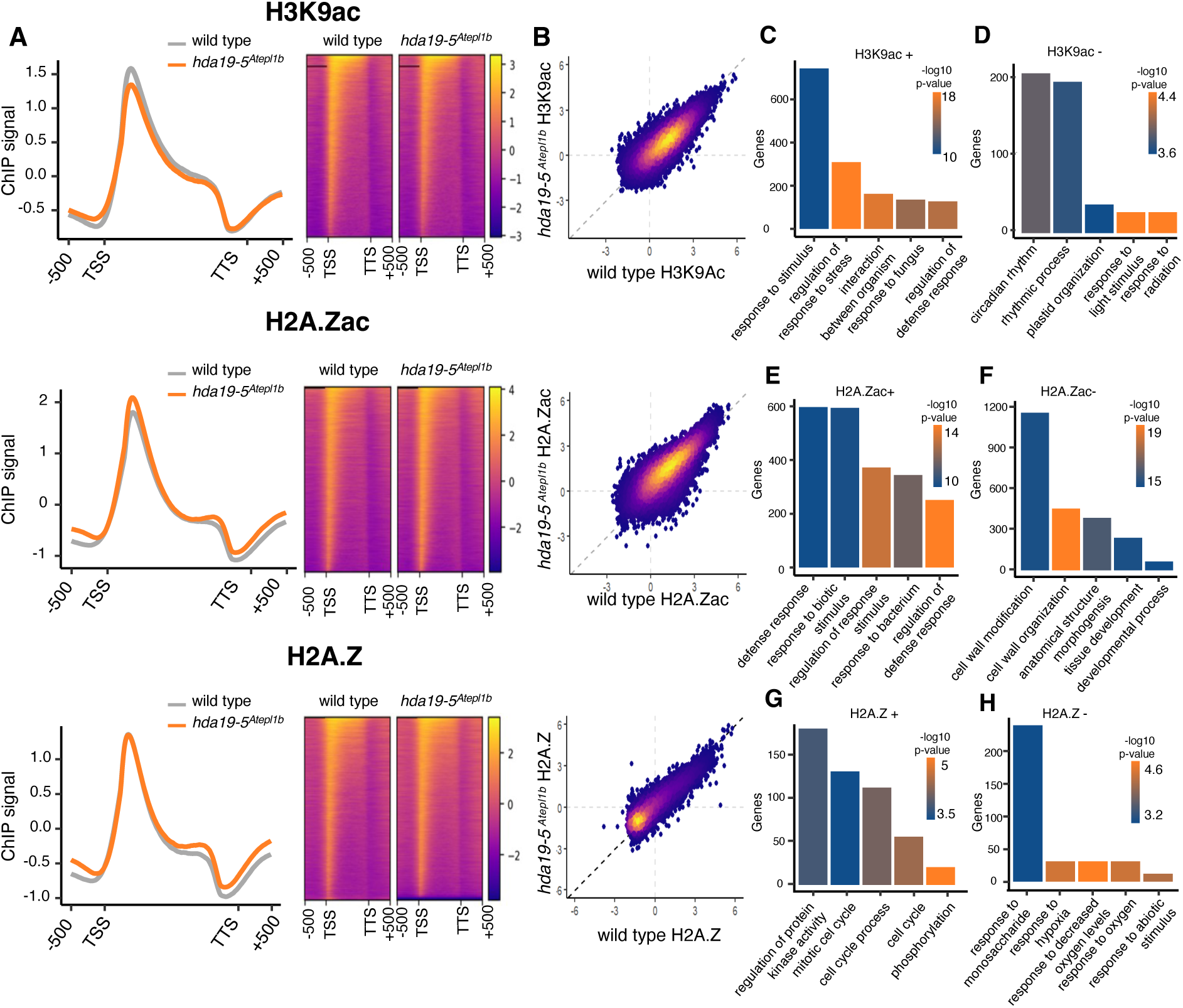
ChIP-seq analysis in hda19-5*^Atepl1b^*. **(A)** Metagene profiles of H3K9ac signal for expressed genes (n = 19,256) in wild-type (WT) plants grown at 22°C (grey) and *hda19-5****^Atepl1b^*** (left). TSS marks the transcription start site and TTS the transcription termination site. Right, heatmaps showing H3K9ac occupancy over expressed genes in WT and *hda19-5****^Atepl1b^***. Colour encodes H3K9ac enrichment over input. **(B)** Scatterplot showing H3K9ac levels at individual genes in WT and *hda19-5****^Atepl1b^***. Each point represents one gene. Colours indicate the amount of H2A.Z in the gene body region. **(C–H)** Gene Ontology (GO) enrichment for genes showing differential enrichment of H3K9ac (top; C–D), H2A.Zac (middle; E–F), and total H2A.Z (bottom; G–H) in *hda19-5****^Atepl1b^*** compared with WT. The y-axis indicates the number of genes in each enriched category, and the x-axis shows GO terms. Bar colour reflects statistical significance (−log₁₀ FDR-adjusted *p*-value). For each mark, the five most representative and abundant GO categories are shown.

At the level of individual genes, however, we observed pronounced heterogeneity: some loci showed increased acetylation in *hda19-5^Atepl1b^*, whereas others displayed reduced signal (Fig. 3B). This prompted us to analyse H3K9ac, H2A.Zac, and H2A.Z enrichment in functionally defined gene subsets. Using differential occupancy analysis (log₂FC > 0.6 or log₂FC < −0.6; Shao et al., 2012), we identified 1422 genes with elevated H3K9ac in *hda19-5^Atepl1b^*(Fig. 3C). Gene Ontology analysis revealed strong enrichment of stress-related categories among these hyperacetylated genes (Fig. 3C; Supplementary Fig. 20A; Supplementary Tables 10–11). Conversely, 1642 genes showed reduced H3K9ac levels, and these were predominantly associated with responses to light and radiation (Fig. 3D; Supplementary Table 11). Notably, GO enrichment for hyperacetylated genes was substantially more significant than for genes with reduced H3K9ac (Supplementary Table 11).

Changes in H2A.Z acetylation were even more widespread than those in H3K9ac. In *hda19-5^Atepl1b^*, 4204 genes showed increased H2A.Zac signal, many of which were annotated with defence-related GO terms (Fig. 3E; Supplementary Fig. 20B; Supplementary Table 11). In parallel, 4357 genes exhibited reduced H2A.Zac levels, and these were enriched for developmental processes (Fig. 3F; Supplementary Table 11). Together, these results suggest that HDA19 catalyses H2A.Z deacetylation at stress-related genes, while its loss is associated with diminished H2A.Z acetylation at developmental genes, possibly reflecting reduced histone acetyltransferase activity at these loci.

We next assessed differential occupancy of total H2A.Z in *hda19-5^Atepl1b^*. We detected 3489 genes with increased H2A.Z levels; however, GO categories and associated *p*-values for these genes were relatively modest (Fig. 3G; Supplementary Table 11), with some enrichment in cell cycle–related terms. In contrast, only 822 genes showed reduced H2A.Z occupancy, with the largest group belonging to the “response to abiotic stimulus” category (Fig. 3H; Supplementary Table 11). These findings suggest that changes in total H2A.Z levels may largely reflect indirect consequences of altered histone acetylation in the absence of HDA19, potentially influencing the activity of other complexes such as SWR1 (Nie et al., 2019).

Consistent with this notion, inspection of H2A.Z profiles around the TSS of genes with elevated H2A.Zac revealed reduced total H2A.Z occupancy, whereas genes with decreased H2A.Zac showed increased H2A.Z enrichment. A control set of 2500 randomly selected genes displayed intermediate levels. These patterns indicate that H2A.Z acetylation promotes H2A.Z eviction or turnover from chromatin, supporting its role as a dynamic regulatory modification rather than a stabilizing mark (Supplementary Fig. 20C).

### Stress gene expression in *hda19-5^Atepl1b^* is shaped by H3K9ac and H2A.Zac changes

To assess how H3K9 and H2A.Z acetylation affect gene expression, we examined these chromatin marks at the 656 genes upregulated in *hda19-5^Atepl1b^*(Fig. 2A–B). Compared with wild type, *hda19-5^Atepl1b^* showed significantly higher levels of both H3K9ac and H2A.Zac at these loci (two-sided unpaired Wilcoxon test, *p* = 0.02 and *p* = 0.03, respectively; Fig. 4A). By contrast, total H2A.Z levels at these genes did not differ between wild type and *hda19-5^Atepl1b^* (*p* = 0.86; Fig. 4A–B). These data suggest that increased histone acetylation in *hda19-5^Atepl1b^* contributes to the upregulation of a subset of stress-response genes (Fig. 2C). Notably, gene-body H2A.Z levels (gbH2A.Z; 1250–2000 bp downstream of the TSS) remained unchanged upon transcriptional activation, consistent with ongoing H2A.Z incorporation by the SWR1 complex following initial eviction (Krall and Deal, 2024).

**Fig. 4.**
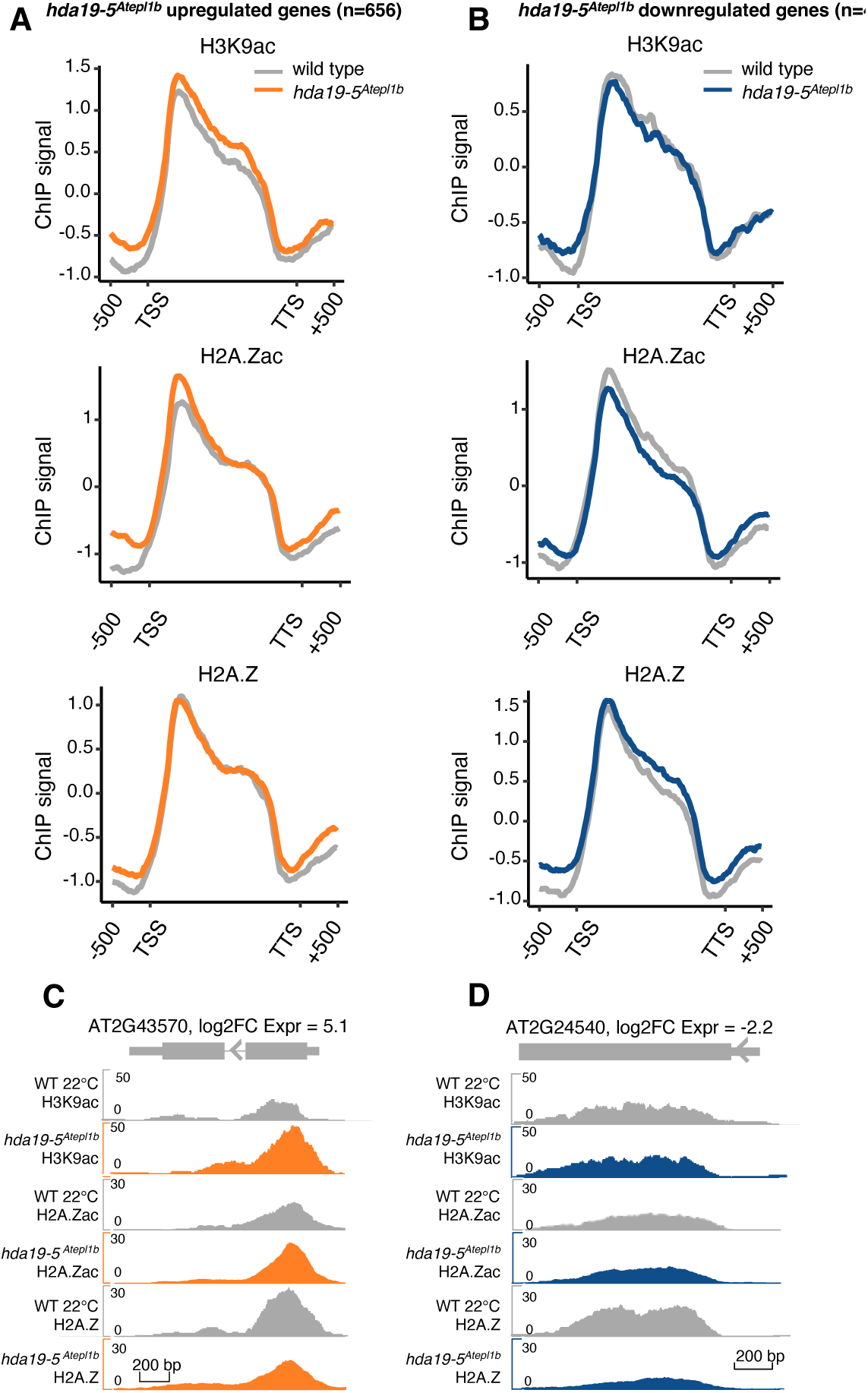
Levels of chromatin modifications at genes dysregulated in *hda19-5^Atepl1b^*. **(A)** Metagene profiles of H3K9ac, H2A.Zac, and total H2A.Z at the 656 genes upregulated in *hda19-5****^Atepl1b^***. Orange lines indicate signal in *hda19-5****^Atepl1b^***; grey lines indicate wild-type signal. **(B)** Metagene profiles of H3K9ac, H2A.Zac, and total H2A.Z at the 405 genes downregulated in *hda19-5****^Atepl1b^***. Blue lines indicate signal in *hda19-5****^Atepl1b^***; grey lines indicate wild-type signal. **(C)** Genome browser snapshot showing increased H3K9ac and H2A.Zac at the +1 nucleosome of a representative stress-related gene upregulated in *hda19-5****^Atepl1b^***. **(D)** Genome browser snapshot showing changes in H3K9ac, H2A.Zac, and H2A.Z at a representative photosynthesis-related gene downregulated in *hda19-5****^Atepl1b^***.

We next analysed histone marks at the 405 genes downregulated in *hda19-5^Atepl1b^*. H3K9ac levels at these loci did not differ significantly between genotypes (two-sided unpaired Wilcoxon test, *p* = 0.26). In contrast, H2A.Zac levels were reduced in *hda19-5^Atepl1b^* (*p* = 0.005), while gbH2A.Z levels were increased (*p* = 0.007; Fig. 4B). The combination of decreased H2A.Z acetylation and elevated gene-body H2A.Z occupancy may contribute to repression of these genes in *hda19-5^Atepl1b^* (Fig. 4C–D).

Taken together, our ChIP-seq data indicate that, despite extensive bidirectional changes in histone acetylation (both hyper- and hypoacetylation), the *hda19-5^Atepl1b^*phenotype – characterized by elevated expression of stress-response genes – is driven predominantly by increased H3K9 and H2A.Z acetylation, with relatively minor contributions from changes in total H2A.Z occupancy.

### Increased temperature modifies H3K9ac and H2A.Zac at stress-related genes in wild-type plants, resembling changes in the *hda19-5^Atepl1b^* mutant

To assess how H3K9ac and H2A.Zac contribute to gene regulation at elevated temperature, we performed ChIP-seq for these marks in wild-type plants grown at 27°C. We first examined H3K9ac, H2A.Z, and H2A.Zac profiles across expressed genes (the same set as in Fig. 3; n = 19,256). In wild-type plants at 27°C, global H3K9ac levels were not significantly altered compared with 22°C (two-sided unpaired Wilcoxon test, *p* = 0.11; Supplementary Fig. 21A). By contrast, we detected a modest increase in H2A.Zac at the +1 nucleosome (*p* = 0.003). Consistent with previous reports of H2A.Z eviction at higher temperatures (Kumar and Wigge, 2010; van der Woude et al., 2019), total H2A.Z levels were reduced over gene bodies (*p* = 1.32 × 10⁻¹¹). Importantly, at 27°C individual genes showed substantial, locus-specific changes in histone modifications, with the most pronounced differences observed for H2A.Z acetylation (Supplementary Fig. 21B).

We next identified genes displaying differential enrichment of H3K9ac, H2A.Zac, and total H2A.Z in wild-type plants at 27°C relative to 22°C (Supplementary Fig. 21C–D). GO analysis revealed that genes with increased H3K9ac (n = 1,639) and increased H2A.Zac (n = 5,015) were enriched for stress-response categories, as were genes with the strongest depletion of gene-body H2A.Z (n = 1,083) (Supplementary Tables 10 and 12). Notably, the GO terms associated with hyper- and hypoacetylated genes at 27°C closely resembled those identified in *hda19-5^Atepl1b^* (Fig. 3C–D; Supplementary Tables 11–12).

To directly compare these responses, we analysed overlaps between genes differentially enriched for H3K9ac, H2A.Zac, and total H2A.Z in *hda19-5^Atepl1b^*at 22°C and in wild type at 27°C, both relative to wild type at 22°C. All six types of chromatin changes showed significant overlap, but the largest was observed for H2A.Z hyper-and hypoacetylation (77.9% and 85.4% shared genes, respectively; Fig. 5A, C). H2A.Z acetylation also affected the greatest number of common targets, with more than 3,000 genes showing significant increases or decreases in H2A.Zac in both contexts (Fig. 5A, C).

**Fig. 5.**
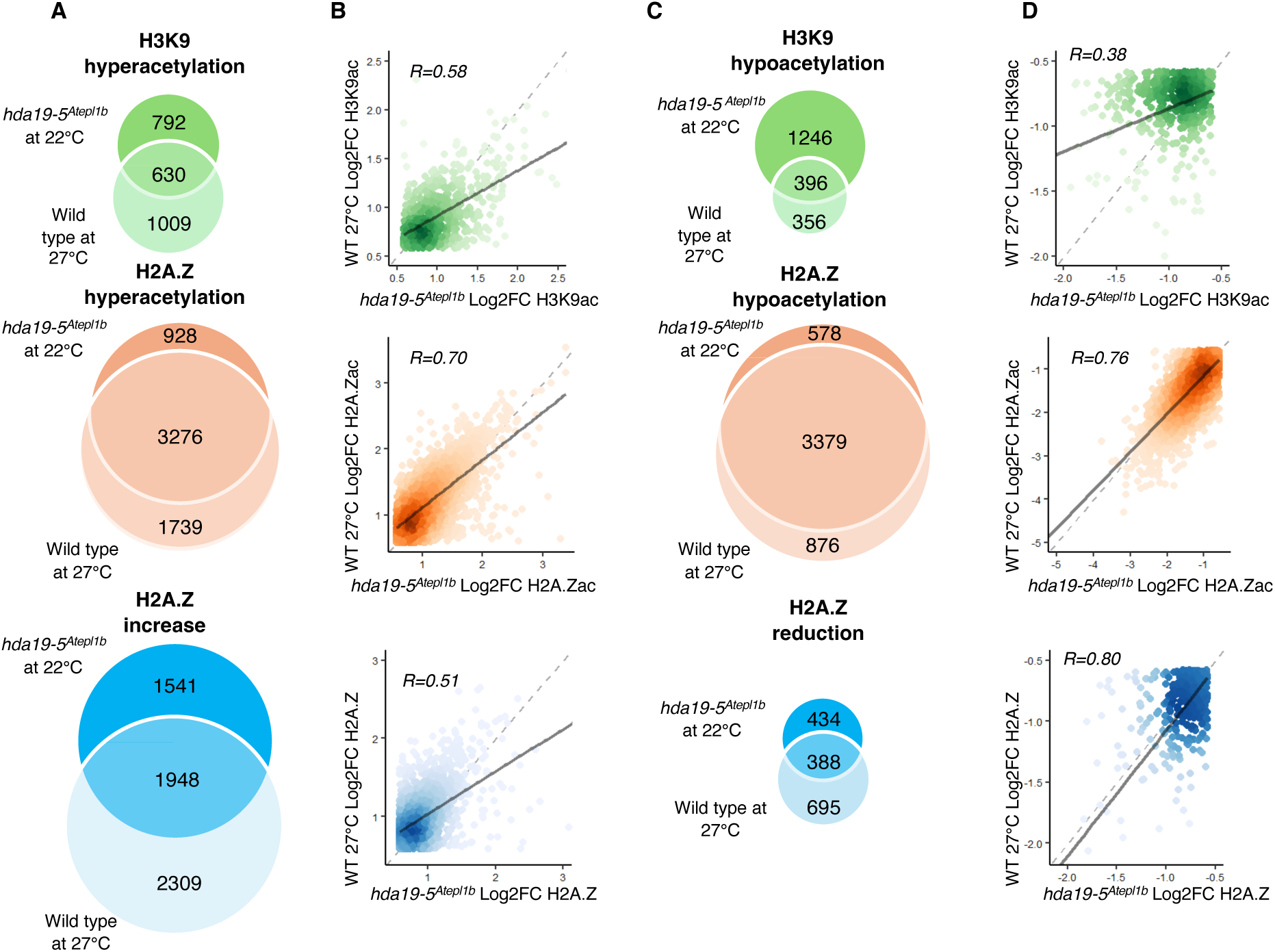
HDA19 and NuA4 co-regulate H2A.Z acetylation. **(A)** Venn diagrams showing overlaps between genes with increased H3K9ac, H2A.Zac, or total H2A.Z in *hda19-5****^Atepl1b^*** at 22°C and wild-type plants grown at 27°C, each compared with WT at 22°C. **(B)** Scatter plots showing fold changes of H3K9ac, H2A.Zac, and total H2A.Z for genes commonly gaining each mark in *hda19-5****^Atepl1b^*** and wild type at 27°C. Pearson correlation coefficients and associated *p*-values (*p* < 2.2 × 10⁻¹⁶ for all marks) are indicated. **(C)** Venn diagrams showing overlaps between genes with decreased H3K9ac, H2A.Zac, or total H2A.Z in *hda19-5****^Atepl1b^*** at 22°C and WT at 27°C, each compared with WT at 22°C. **(D)** Scatter plots showing fold changes of H3K9ac, H2A.Zac, and total H2A.Z for genes commonly losing each mark in *hda19-5****^Atepl1b^*** and wild type at 27°C. Pearson correlation coefficients and associated *p*-values (*p* < 2.2 × 10⁻¹⁶ for all marks) are indicated.

We then asked whether genes with similar patterns of hyper- or hypoacetylation, or with gains or losses in total H2A.Z, exhibit stronger chromatin changes in *hda19-5^Atepl1b^* than in wild type at 27°C. Differentially enriched genes showed comparable changes in H3K9ac and H2A.Zac in both backgrounds, with no systematic increase in effect size in *hda19-5^Atepl1b^*(Fig. 5B, D). Once again, H2A.Z acetylation changes were the most strongly correlated between the two conditions (R = 0.70 for hyperacetylated and R = 0.76 for hypoacetylated genes). For most genes, fold changes in acetylation and H2A.Z levels were moderate.

Taken together with the similar patterns of gene expression dysregulation in wild type at 27°C and *hda19-5^Atepl1b^* at 22°C (Fig. 2B), these data provide a mechanistic explanation for why *hda19-5^Atepl1b^*phenocopies wild-type plants grown at elevated temperature. We propose that the absence of HDA19 “primes” stress-related genes by preventing deacetylation of H3 and H2A.Z, thereby enabling their activation in the absence of an external thermal stimulus.

### Convergent regulation of stress genes by NuA4 and HDA19

Unlike in yeast, where loss of specific histone deacetylases can partially rescue NuA4 deficiency (Lin et al., 2008), *Arabidopsis* HDAC mutations did not suppress the *Atepl1-1* phenotype. On the contrary, additional *hdac* mutations tended to aggravate it (Supplementary Fig. 14). Because *Atepl1-1* shows broad transcriptional activation of stress-related genes (Bieluszewski et al., 2022a), we asked whether HDA19 co-regulates a subset of *Atepl1*-upregulated genes. Strikingly, more than 75% (501) of the genes upregulated in *hda19-5^Atepl1b^* were also upregulated in *Atepl1-2* (a CRISPR-derived *Atepl1* null allele) (Fig. 6A), despite lack of any accumulated effect of *Atepl1b-1* on plant development or biochemistry (Supplementary Fig. 15-17, Fig. 3). For most of these 501 genes, the magnitude of transcriptional activation was higher in *Atepl1-2* than in *hda19-5^Atepl1b^* (Fig. 6B). This extensive overlap suggests that the majority of *hda19-5^Atepl1b^*-upregulated genes are controlled by NuA4-dependent mechanisms, likely involving H2A.Z deposition and/or acetylation.

**Fig. 6.**
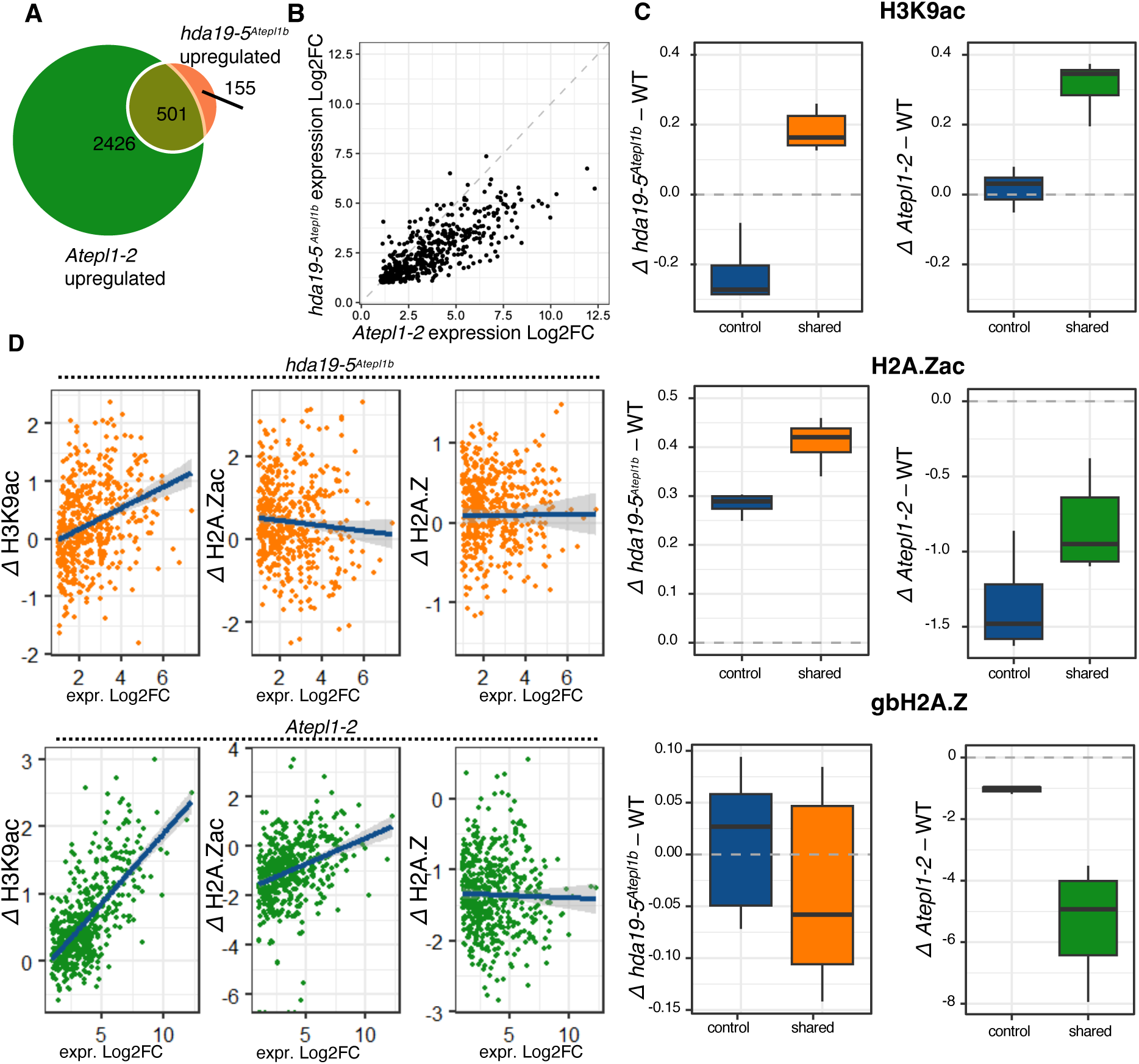
NuA4 and HDA19 cooperate to repress stress-related genes. **(A)** Overlap between genes upregulated in *hda19-5****^Atepl1b^*** and *Atepl1-2*. Statistical significance of the overlap was assessed using a hypergeometric test (*p* < 2.2 × 10⁻¹⁶). **(B)** Scatter plot showing the correlation of expression fold changes for the 501 genes co-upregulated in *hda19-5****^Atepl1b^*** and *Atepl1-2*. Each point represents one gene. **(C)** Boxplots showing H3K9ac, H2A.Zac, and total H2A.Z levels for 2500 control expressed genes (control, blue) and for the 501 co-upregulated (shared) genes. Left, comparison between wild type (WT) and *hda19-5****^Atepl1b^*** (shared genes in orange); right, comparison between WT and *Atepl1-2* (shared genes in green). **(D)** Correlations between changes in H3K9ac, H2A.Zac, or total H2A.Z and expression fold change for the 501 co-upregulated genes. Pearson correlation coefficients and *p*-values are shown for each modification.

To explore this possibility, we analysed ChIP-seq data for H3K9ac, total H2A.Z, and H2A.Zac in wild type and *hda19-5^Atepl1b^*, together with previously published data for *Atepl1-2* (Bieluszewski et al., 2022a). We focused on histone modification levels at the 501 genes co-upregulated in *hda19-5^Atepl1b^* and *Atepl1-2*, and compared them with 2500 control genes.

In *hda19-5^Atepl1b^*, these 501 shared genes exhibited significantly higher H3K9ac and H2A.Zac levels than the control set (two-sided unpaired Wilcoxon test, *p* = 4.0 × 10⁻¹⁵ and *p* = 5.8 × 10⁻¹¹, respectively; Fig. 6C). In contrast, total H2A.Z levels showed a small but significant decrease compared with control genes (*p* = 1.57 × 10⁻⁵).

In the *Atepl1-2* background, H3K9ac levels over the 501 shared genes were slightly higher than in *hda19-5^Atepl1b^* (*p* = 4.0 × 10⁻¹⁵; Fig. 6C). By contrast, both H2A.Zac and gene-body H2A.Z (gbH2A.Z) were strongly reduced in *Atepl1-2* for both control and co-upregulated genes compared with wild type and *hda19-5^Atepl1b^*, reflecting the global impact of NuA4 loss on H2A.Z and its acetylated form (*p* = 1.1 × 10⁻⁸ and *p* = 2.2 × 10⁻¹⁶, respectively). Notably, the reduction in H2A.Zac at the 501 co-upregulated genes was less pronounced than at control genes, suggesting that these stress-related genes normally maintain relatively low basal H2A.Zac levels (Fig. 6C).

We then examined how the same 501 genes behave in wild-type plants grown at elevated temperature (27°C; Supplementary Fig. 22). In this context, H3K9ac levels increased significantly (*p* = 4.0 × 10⁻¹⁵), whereas H2A.Zac levels did not differ from those in the control gene set (*p* = 1). Total H2A.Z was slightly more reduced in the 501-gene group than in controls (*p* = 0.001). These observations support the idea that the acetylation status of H3K9 and H2A.Z at a subset of stress genes – under joint control of HDA19 and NuA4 – modulates their transcriptional activity under non-inducing conditions.

Although co-upregulated genes show increased acetylation in both *hda19-5^Atepl1b^*and *Atepl1-2*, they are more strongly transcriptionally activated in *Atepl1-2* (Fig. 6B–C). To test whether this difference can be explained by distinct chromatin landscapes, we correlated expression fold changes with changes in H3K9ac, H2A.Zac, and H2A.Z at individual genes (Fig. 6D). Gene expression changes correlated positively with H3K9ac gains in both *hda19-5^Atepl1b^* and *Atepl1-2* (Pearson’s r = 0.32, *p* = 6.5 × 10⁻¹³; and r = 0.67, *p* < 2.2 × 10⁻¹⁶, respectively). In *hda19-5^Atepl1b^*, however, we detected no significant correlation between expression and changes in H2A.Zac (r = −0.08, *p* = 0.08) or total H2A.Z (r = 0.007, *p* = 0.87). By contrast, in *Atepl1-2* the expression fold change correlated with H2A.Zac (r = 0.34, *p* = 2.52 × 10⁻¹⁵), but not with depletion of H2A.Z (r = −0.002, *p* = 0.58).

These results suggest that H3K9 acetylation plays a stronger role than H2A.Zac in controlling the expression of the 501 commonly regulated genes in *hda19-5^Atepl1b^*. H2A.Zac may act as an accessory modification that is not strictly required for transcriptional activation, consistent with its behaviour in *Atepl1-2* mutants. We conclude that the stronger transcriptional activation observed in *Atepl1-2* reflects the combined effect of a greater gain in H3K9ac and a pronounced depletion of gbH2A.Z, which normally contributes to repression at these stress-related loci.

### Loss of HDA19 leads to increased H2A.Z acetylation near its binding sites

To further clarify the role of H2A.Zac in transcriptional activation and strengthen the functional link between HDA19 and H2A.Z deacetylation, we compared a recently published ChIP-seq dataset of HDA19 occupancy in the *Arabidopsis* genome with our own data (Liu et al., 2025). We first selected well-defined H2A.Z peaks (width < 500 bp) identified in *hda19-5^Atepl1b^* grown at 22°C and intersected these regions with HDA19 binding sites. This analysis identified 6,678 H2A.Z peaks overlapping HDA19-bound regions and 9,686 H2A.Z peaks lacking HDA19 occupancy.

We then quantified changes in H2A.Z acetylation in *hda19-5^Atepl1b^*relative to wild type at these two groups of peaks. H2A.Z peaks co-occupied by HDA19 showed markedly higher H2A.Zac levels in *hda19-5^Atepl1b^* compared with H2A.Z peaks without HDA19 binding (Fig. 7A–B). Notably, H2A.Z/HDA19-overlapping peaks were frequently associated with genes involved in essential biological processes and stimulus responses (Fig. 7C; Supplementary Table 13). In contrast, H2A.Z peaks lacking HDA19 did not show significant GO term enrichment, but instead were under-represented for various response-related categories (Supplementary Table 13). These observations support a model in which HDA19 locally reduces H2A.Z acetylation at H2A.Z-enriched regulatory regions.

**Fig. 7.**
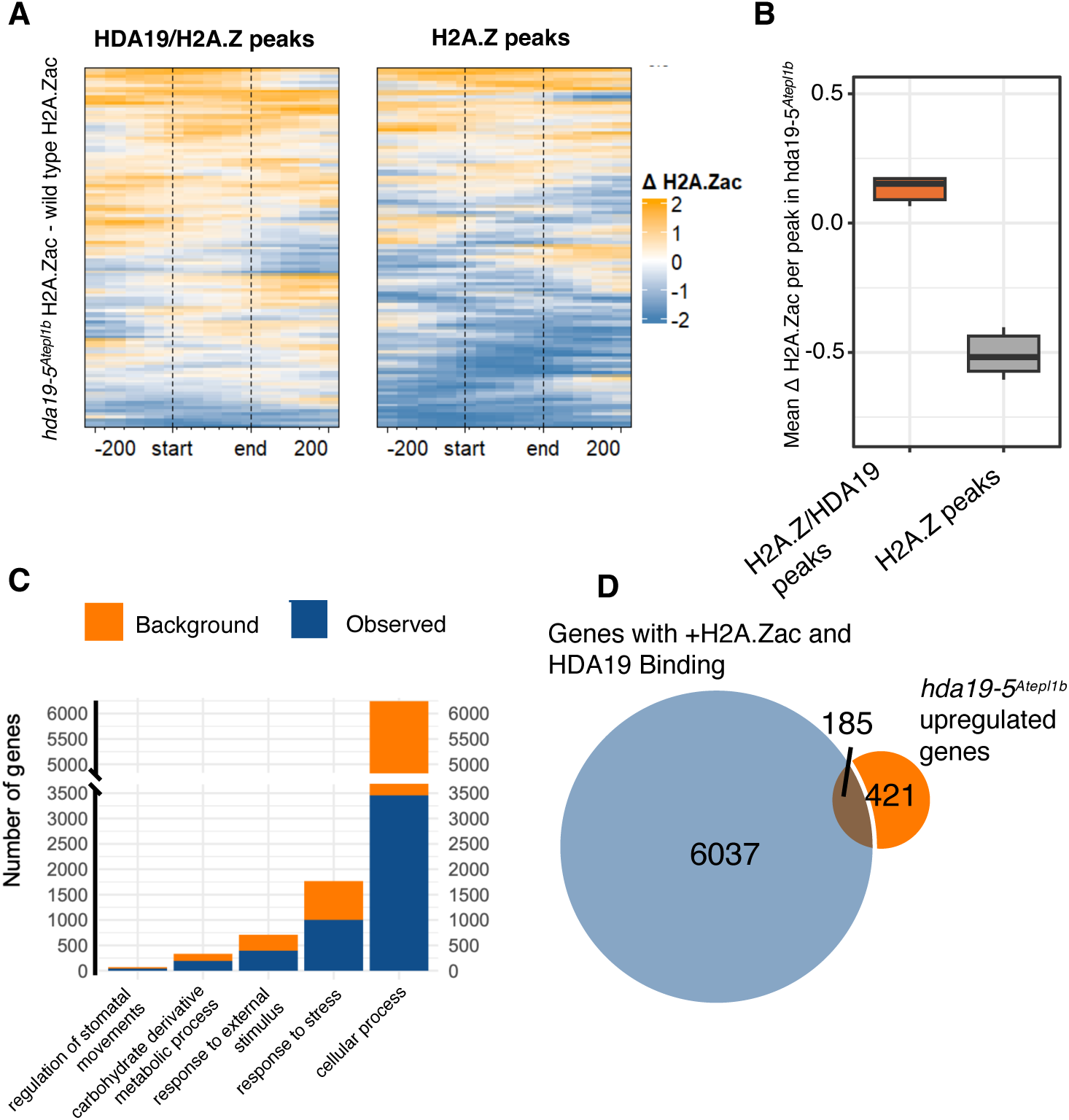
H2AZac is increased around HDA19 binding sites in *hda19-5^Atepl1b^.* **(A)** Heatmaps showing differences in H2A.Zac signal between wild type (WT) and *hda19-5****^Atepl1b^*** around H2A.Z peaks that are co-bound by HDA19 (left) or bound only by H2A.Z (right). **(B)** Mean H2A.Zac difference between WT and *hda19-5****^Atepl1b^*** for the two groups of peaks shown in (A). **(C)** Gene ontology (GO) enrichment analysis for genes associated with HDA19/H2A.Z co-bound peaks. The top five non-redundant GO terms were selected using *rrvgo* (Sayols, 2023). ‘Background’ shows all the genes corresponding to a given GO term, while ‘Observed’ shows genes co-bound by H2A.Z and HDA19. **(D)** Venn diagram showing the overlap between genes upregulated in *hda19-5****^Atepl1b^*** and genes co-occupied by HDA19 and H2A.Z.

To examine how elevated H2A.Zac levels affect transcription, we compared genes upregulated in *hda19-5^Atepl1b^* (Supplementary Table 13) with genes bound by HDA19 that also displayed higher H2A.Zac signal in the mutant. The overlap between these two sets was moderate (Fig. 7D), indicating that increased H2A.Zac alone is not sufficient and not strictly required for transcriptional activation. This conclusion is consistent with our earlier analyses in *Atepl1-2* and *hda19-5^Atepl1b^*, which showed that changes in H3K9ac correlate strongly with gene activation, highlighting H3K9 acetylation as the primary determinant of transcriptional output, while H2A.Zac functions as a modulatory mark controlled by the HDA19/NuA4 axis (Fig. 6D).

## Discussion

Although deacetylation of histones H3 and H4 by HDACs has been extensively studied, deacetylation of the histone variant H2A.Z has received much less attention in plants. In yeast and humans, H2A.Z deacetylases have been implicated in transcriptional regulation at specific loci (Link et al., 2018; Mehta et al., 2010). In plants, H2A.Z has been studied primarily in the context of heat responses, where it has been linked to chromatin accessibility and transcriptional activation (León et al., 2024; Tasset et al., 2018; Wollmann et al., 2017; Sura et al., 2017). Here, we provide evidence for a functional connection between HDA19, H2A.Z, and heat-related stress responses.

We found that *hda19-5^Atepl1b^* mutants exhibit pronounced developmental and fertility defects, which are likely caused, at least in part, by altered expression of stress-related genes (Fig. 1). Transcriptomic profiling showed that *hda19-5^Atepl1b^* displays misregulation of biotic stress–responsive genes, but not of canonical heat-responsive genes induced by elevated temperature (Fig. 2). ChIP-seq analysis further revealed that genes involved in environmental responses are hyperacetylated at H3K9 and H2A.Z in *hda19-5^Atepl1b^*, without major changes in gene-body H2A.Z levels (Fig. 3). Upregulated genes in *hda19-5^Atepl1b^* showed increased H3K9ac and H2A.Zac, which likely contributes to their activation in the absence of external stress (Fig. 4). Consistent with this idea, chromatin changes in *hda19-5^Atepl1b^*strongly overlap with those in wild-type plants exposed to heat stress (Fig. 5), suggesting that the hyperacetylation of these loci is a direct consequence of HDA19 loss and functionally mimics physiological thermal cues.

Strikingly, most genes upregulated in *hda19-5^Atepl1b^*are also upregulated in *Atepl1-2*, a mutant lacking a functional NuA4 acetyltransferase and displaying a drastic reduction in both acetylated and non-acetylated H2A.Z. In *Atepl1-2*, H2A.Z and H4 acetylation are depleted, and the activation of stress-responsive genes is primarily driven by compensatory hyperacetylation of H3. By contrast, in *hda19-5^Atepl1b^*total H2A.Z levels remain relatively stable, whereas H2A.Zac is elevated at both stress-induced and transcriptionally unaffected genes. This pattern supports a direct role for HDA19 in H2A.Z deacetylation, rather than H2A.Zac changes being a mere secondary consequence of transcriptional activation. Together, these observations indicate that in both *Atepl1-2* and *hda19-5^Atepl1b^*, H3 acetylation acts as the principal driver of stress gene induction, while H2A.Zac plays a modulatory role (Fig. 6).

Further integration of our data with recently published HDA19 ChIP-seq maps in *Arabidopsis* (Liu et al., 2025) showed that, in *hda19-5^Atepl1b^*, H2A.Z peaks overlapping HDA19 binding sites display higher H2A.Zac levels than H2A.Z peaks not bound by HDA19 (Fig. 7). These HDA19/H2A.Z co-bound peaks are preferentially associated with genes involved in essential biological processes and stimulus responses, reinforcing the notion that HDA19 directly deacetylates H2A.Z at regulatory loci. However, the overlap between genes with elevated H2A.Zac and those upregulated in *hda19-5^Atepl1b^* was only moderate (Fig. 7D), indicating that increased H2A.Zac alone is not sufficient to enhance transcription. This is consistent with recent findings that the N-terminal tail of H2A.Z, which is a major target of acetylation, is largely dispensable for plant development but crucial for proper stress responses (Sijacic et al., 2025). Combined with our results, this suggests that the primary role of H2A.Z acetylation may not be to directly promote transcription, but rather to counteract repressive features of gene-body H2A.Z. This would explain why removal of the H2A.Z tail does not impair basal transcriptional activation but affects the dynamic control of stress responses.

In yeast, mutations in specific HDACs can suppress the defects caused by loss of NuA4 activity (Lin et al., 2008). Similarly, in *Arabidopsis*, loss of the SAGA component GCN5 can be alleviated by *hda19* mutation (Benhamed et al., 2006). In contrast, we observed that combined loss of NuA4 and HDA19 leads to additive or even aggravated phenotypes rather than mutual suppression (Supplementary Fig. 14). This raises two possibilities: either higher-order combinations of HDAC mutations might be required to compensate for NuA4 loss in plants, or there is no alternative acetyltransferase capable of functionally replacing NuA4 under conditions of reduced histone acetylation.

Among chromatin-associated complexes, the PEAT (PWWPs–EPCRs–ARIDs–TRBs) complex has emerged as a key regulator of histone deacetylation in *Arabidopsis* (Tan et al., 2018). PEAT has been implicated in heterochromatin formation through interactions with HDA6 and HDA9. Intriguingly, two PEAT subunits, EPCR1 and EPCR2, share distant homology with the EPL1A and EPL1B subunits of NuA4, and both complexes share HAM1/2 acetyltransferases, whereas the remaining subunits are complex-specific (Tan et al., 2018). A recent study showed that NuA4 and PEAT co-occupy a substantial fraction of transcription start sites, where their cooperation is required for histone acetylation and transcriptional activation (Zheng et al., 2023). Mutations in PEAT components cause pleiotropic growth defects and dysregulation of both stress-responsive and developmental genes (Zheng et al., 2023). Furthermore, the same study demonstrated that the histone deubiquitinase UBP5 interacts with PEAT PWWP subunits, thereby coupling histone acetylation with removal of repressive H2A ubiquitination (Zheng et al., 2023). Previous work has shown that H2A.Z is stabilized within nucleosomes by PRC1-mediated monoubiquitination, which is associated with transcriptional repression (Gómez-Zambrano et al., 2019). Because UBP5 removes ubiquitin from H2A, it will be important to determine whether the PEAT–UBP5 module also targets H2A.Zub at PEAT-bound loci (Godwin et al., 2024).

Taken together, our data support a model in which NuA4 and histone deacetylases, particularly HDA19, cooperate to prevent spurious activation of stress-inducible genes. This cooperation extends beyond simple removal of H2A.Zac, as aberrant accumulation of this mark alone is not sufficient to trigger stress responses. Instead, efficient repression of many stress genes appears to require the presence of H2A.Z in gene bodies together with low H3 acetylation (Fig. 8). Future work should investigate how HDA19 functionally interacts with other chromatin-modifying complexes such as PEAT, whether these regulatory principles are conserved across different environmental conditions and developmental stages, and how the balance between H3 acetylation and H2A.Z-based mechanisms fine-tunes the responsiveness of stress-related loci.

**Fig. 8.**
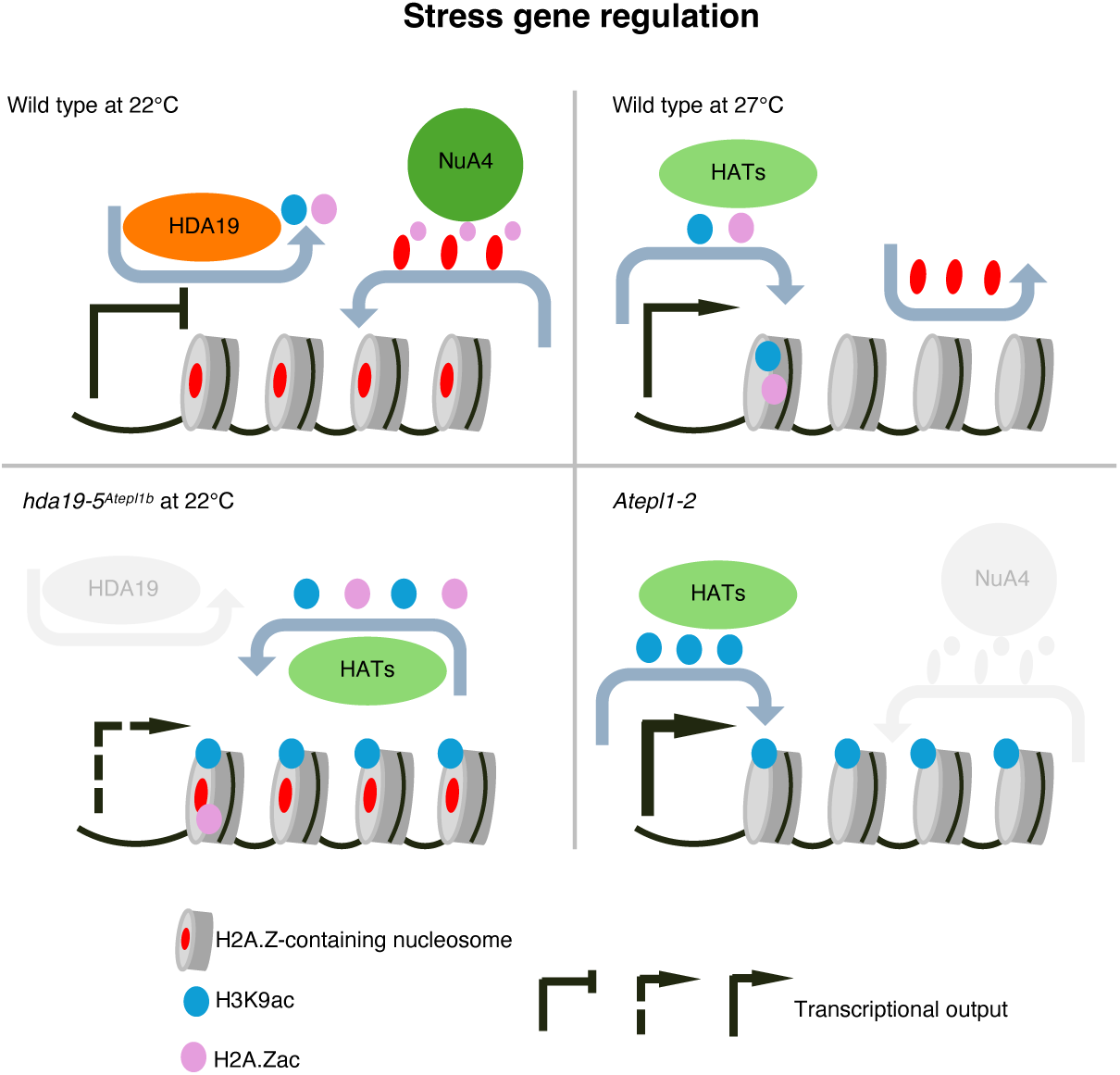
Regulation of stress gene transcription by histone acetylation and H2A.Z. In wild-type plants growing under physiological conditions, chromatin at stress genes is maintained in a repressive state through coordinated action of HDA19 and the NuA4 complex. Upon exposure to elevated temperature, acetyl groups are deposited by histone acetyltransferases (HATs) at +1 nucleosome, combined with removal of gene-body H2A.Z which leads to transcriptional activation of normally repressed genes. In *hda19-5^Atepl1b^* background, the activity of HATs (including NuA4), cannot be balanced by histone deacetylation, leading to increased levels of H3K9ac and H2A.Zac and in turn upregulation of a subset of stress genes. This upregulation however does not span all the stress genes activated in wild-type plants grown at 27°C and is counteracted by H2A.Z occupancy of gene-body regions. In *Atepl1-2* mutant, H4ac (not shown), H2A.Zac and H2A.Z are abolished. Elevated H3K9ac levels, together with H2A.Z depletion, lead to strong transcriptional activation of HDA19-NuA4 target genes.

## Materials and Methods

### Plant growth conditions and material

Plants were grown on pellets in long-day conditions at 22°C or 27°C, at 70% humidity and 80–90 μmol/(m^2^s) PAR. Seeds were stratified at 4°C for 48 hours prior to transferring the pellets to growth chambers. The *arp6-1* mutant was a gift from prof. Kyuha Choi (Choi et al., 2007). T-DNA insertion lines for *Atepl1-1a* and *Atepl1-1b* were obtained from NASC (SAIL_239_D12 and SALK_094941; respectively).

### CRISPR-Cas9 mutagenesis of histone deacetylases

GuideRNA sequences were designed with use of the CRISPOR software (Haeussler et al., 2016). The generated sequences were chosen according to their specificity, efficiency and off-target number. The gRNAs were targeted to induce DSBs within the same exon sequences, usually close to 5’ end of the gene. Genetic constructs were prepared as described previously with use of ClonExpress MultiS One Step Cloning Kit (Vazyme, Cat. No. C113-02) (Bieluszewski et al., 2022b). Columbia-0 (Col-0) plants were transformed by *Agrobacterium*-mediated floral dip (GV3101 strain). Resulting T1 seeds were selected under epifluorescent macroscope. Transformants were PCR-screened for the presence of construct and mutation in the targeted histone deacetylase with Phire Plant Direct PCR Master Mix (Thermo Fisher Scientific, Cat. No. F160L). Propagation of T2 plants was based on the presence of mutation and Cas9 vector was segregated out by selecting non-fluorescent seeds. The deletions found in *hdac* mutants were confirmed with Sanger sequencing. Oligonucleotide sequences used in the experiments are listed in Supplementary Table 14.

### Phenotypic analysis

T3 construct-free *hdac* mutants were grown for 28 days in conditions described above. Representative rosettes were collected and taped into paper sheet, which was later scanned. The measurements of the rosette diameter was carried out in the ImageJ software and Microsoft Excel. Flowering time was determined on the number of primary leaves when the plant developed fully open first flowers. The collected siliques were stored in 80% ethanol for 2 days at 4°C and later photographed under the macroscope. First 10 siliques were omitted from fertility tests.

### Assessment of pollen viability and density

Pollen viability and density were assessed using Alexander staining (10% ethanol, 0.01% Malachite green, 25% glycerol, 0.05% Fuchsin acid, 0.005% Orange G and 4% glacial acetic acid), as detailed in Alexander (Alexander, 1969) and Hord et al. (Hord et al., 2008).

For pollen viability, 1500 pollen grains collected from open flowers of three plants per genotype were analysed. Briefly, mature flowers were immersed in a drop of Alexander staining on a slide and viability of pollen grains was evaluated at 10X magnification under a Leica DM4 B using the bright field. Viable pollen grains are coloured in magenta/red and perfectly round shaped while the non-viable ones are green/brown and lost their round shape.

For pollen density, the number of viable pollen grains was assessed per anthers of stage 12 flower buds collected from three different plants for each genotype. Briefly, buds were first discoloured using Carnoy fixative (60% ethanol, 30% chloroform, 10% glacial acetic acid) for 1 h and then incubated at 55°C in Alexander staining for at least 7 h. Four to six anthers per flower bud were then dissected and mounted between a slide and cover slip using 10% glycerol. Finally, the sampled anthers were observed under the bright field using a Leica DM4 B at 20X magnification.

### Cytology techniques

Chromosome spreading was performed from at least three biological replicates for every genotype as detailed previously with minor modifications (Ross et al., 1996). Briefly, inflorescences were fixed in Carnoy’s solution (3:1, ethanol: acetic acid) for 24 hours. Buds of ∼0.5 µm were incubated for 2 hours at 37°C in an enzymatic solution of 1X citrate buffer at pH=4.5, with 0.3% Cellulase Onozuka R-10, 0.3% Pectolyase Y23 and 0.3% Cytohelicase. Following two washes in 1X citrate buffer at pH=4.5, buds were then crushed on a slide in a 45% acetic acid solution. The slide was placed for 1 min on a heating plate at 42-45°C all the while spreading the mixture with a needle. Then, Carnoy’s fixative was added around and on the top of the mixture and the slide was washed with the same solution and dried at ambient temperature for 24 hours. Finally, pollen mother cells were stained with DAPI and observed under a Leica DM4 B epifluorescence microscope equipped with a Leica DMC5400 20-megapixel colour CMOS camera, as previously described (Zhu et al., 2021).

### RNA isolation and 3’ RNA-seq

RNA was isolated for *hda19-5* and wild type plants grown in different temperatures with the use of RNeasy Plant Mini Kit (Qiagen, Cat. No. 74904). 250 ng of RNA was used for 3’ RNA-seq library preparation. First, reverse transcription reaction was carried out with barcoded and UMI-containing oligo(dT) primers and SuperScript III (Thermo Fisher Scientific, Cat. No 12574026). cDNA was pooled and next purified with AMPure beads (Beckman Coulter, Cat. No. A63881). A second strand synthesis was performed overnight with the nick translation reaction containing 1x NEBNext Second Strand Synthesis Reaction Buffer (New England Biolabs, Cat. No. B6117S), 1 U RNase H (New England Biolabs, Cat. No. M0297L), 1 U *E*. *coli* DNA ligase I (New England Biolabs, Cat. No. M0205L), 5 U *E*. *coli* DNA polymerase (New England Biolabs, Cat. No. M0209L) and 30 µM dNTPs. The double-stranded cDNA was again purified and tagmented with adapter B-loaded Tn5 transposase. Finally, libraries have been amplified with Q5® High-Fidelity 2X Master Mix (New England Biolabs, Cat. No. M0492S) for 9 cycles, and further sequenced on NovaSeq 6000 in 2 × 100 bp PE mode. 4 samples per condition were analysed (16 total).

In our sequencing strategy, read R1 contained UMI and barcode with low-quality sequences after the run. Read R1 was trimmed to keep only UMI and barcode. Fastq files for read R2 from the pools were demultiplexed into separate fastq files for each replicate using BRBseqTools (Alpern et al., 2019) (v 1.6) Demultiplex with parameters -p UB -UMI 14 -n 1. Read R2 of each library was trimmed to remove potential contamination with poly(A) tail using BRBseqTools (v 1.6) Trim and parameters - polyA 10 -minLength 30. Then mapped using STAR (v 2.7.8a) with parameters --sjdbOverhang 99 --outSAMtype BAM SortedByCoordinate --outFilterMultimapNmax 1 (Dobin et al., 2013). Finally, the deduplicated counts for each gene were obtained using BRBseqTools (v 1.6) CreateDGEMatrix with parameters -p UB -UMI 14 -s yes. Counts were used for differential gene expression analysis using the DESeq2 R package (Love et al., 2014). The Gene Ontology analysis was carried out with use of PantherDB after removal of redundant terms with Revigo (Mi et al., 2013; Supek et al., 2011).

### Chromatin isolation and ChIP-seq

The nuclei were isolated from 3 g of cross-linked fresh tissue per sample with use of Honda Buffer (440 mM sucrose, 25 mM Tris-HCl pH 8.0, 10 mM MgCl_2_, 0.5% Triton X-100, 10 mM β-ME, Roche Protein Inhibitor Cocktail, 10 mM PMSF, 40 mM spermine). Nuclei Lysis Buffer (50 mM Tris-HCl pH 8.0, 10 mM EDTA, 1.6% SDS, Roche Protein Inhibitor Cocktail, 50 nM PMSF) was used to extract chromatin, which was later sonicated for 25 cycles (Diagenode Bioruptor). 100 µL of chromatin was later pre-cleared for 1 h at 4°C with Dynabeads Protein A (Thermo Fisher Scientific, Cat. No. 10002D). Next, chromatin was incubated with 5 µg of selected antibody (α-H3K9ac Millipore, Cat. No. 07352, α-HTA9 Agrisera, Cat. No. AS10718, α-H2A.Zac Diagenode, Cat. No. C15419173) overnight at 4°C. 25 µL of Dynabeads Protein A (Thermo Fisher Scientific, Cat. No. 10002D) were added to the samples and incubated for another hour at 4°C. Beads were washed twice with low-salt (150 mM NaCl), high-salt (500 mM NaCl), LiCl (10 mM Tris-HCl, pH 8.0, 1 mM EDTA, 0.25 M LiCl, 1% Nonidet P-40, 0.5% sodium deoxycholate, 1 mM PMSF) and TE buffers. Chromatin was eluted from the beads through incubation with 0.8 M NaHCO_3_ and 1% SDS in 65°C. Following Proteinase K treatment and phenol/chloroform extraction, DNA was precipitated and 2 ng was used to prepare DNA libraries with MicroPlex Library Preparation Kit v2 (10-13 cycles, Diagenode Cat. No. C05010012). Alongside, 10 ng of Input DNA was used to prepare control libraries (6 cycles of amplification). Sequencing has been performed in 2 x 150 PE NovaSeq 6000 mode.

### ChIP-seq data analysis

Paired-end libraries were demultiplexed and trimmed with fastp (Chen et al., 2018). Paired end libraries were mapped to TAIR10 reference genome with bowtie2 (Langmea and Salzberg, 2013). After obtaining the sorted bam files, duplicate reads were removed. Replicate .bam files were merged and transformed into BigWig for heatmaps and gene metaprofiles, which were prepared with bamCompare and deepTools (Ramírez et al., 2014). Then MAnorm was used to determine differentially occupied regions on input-normalized peak file (Shao et al., 2012). Finally, GO analysis has been carried out with ChIPpeakAnno R package, to determine groups of genes affected by differences in H3K9ac, H2A.Z and H2A.Zac (Zhu et al., 2010). GO terms were filtered with use of Revigo and rrvgo (Supek et al., 2011; Sayols, 2023). 2-3 libraries for each tested antibody were used in the analyses. For visualization, MACS2 software was used to call peaks enriched in comparison to chromatin input libraries with --SPMR --keep-dup all --g 120000000 --extsize 147 --nomodel parameters (Zhang et al., 2008).

### ChIP-qPCR

Nuclei were isolated from 500-700 mg of *Arabidopsis* leaves with use of Honda buffer (440 mM sucrose, 25 mM Tris-HCl pH 8.0, 10 mM MgCl_2_, 0.5% Triton X-100, 10 mM β-ME, Roche Protein Inhibitor Cocktail, 10 mM PMSF, 40 mM spermine). Nuclei Lysis Buffer (50 mM Tris-HCl pH 8.0, 10 mM EDTA, 1.6% SDS, Roche Protein Inhibitor Cocktail, 50 nM PMSF) was used to extract chromatin, which was later sonicated for 25 cycles (Diagenode Bioruptor). 100 µL of chromatin was later pre-cleared for 1 h at 4°C with Dynabeads Protein A (Thermo Fisher Scientific, Cat. No. 10002D) and later immunoprecipitation was carried out through overnight incubation at 4°C with 2 ng of selected antibodies (α-H3 ab1791 Abcam, α-H3K9ac Millipore, Cat. No. 07352, α-H2A.Zac Diagenode Cat. No. C15419173). To capture immunocomplexes, Dynabeads Protein A (Thermo Fisher Scientific, Cat. No. 10002D) were added and incubated for additional 1 h at 4°C. Then, beads were washed twice with low-salt (150 mM NaCl), high-salt (500 mM NaCl), LiCl (10 mM Tris-HCl, pH 8.0, 1 mM EDTA, 0.25 M LiCl, 1% Nonidet P-40, 0.5% sodium deoxycholate, 1 mM PMSF) and TE buffers. Finally, complexes have been eluted by incubation with 10% Chelex. Following treatment with proteinase K, the resulting ChIP-DNA was diluted and used as a template in qPCR reactions with use of SYBR Green (Thermo Fisher Scientific, Cat. No. K0222). Percent of input for each tested histone modifications was normalized to Input DNA and H3 levels. 2 or 3 replicates were used for analysis.

## Data availability

Raw read files for 3’RNA-seq and ChIP-seq have been deposited in the GEO database under GSE275206 (https://www.ncbi.nlm.nih.gov/geo/query/acc.cgi?acc=GSE275206 reviewer password: **qzgfqoiultgjhkn)** and GSE275207 (https://www.ncbi.nlm.nih.gov/geo/query/acc.cgi?acc=GSE275207 reviewer password: **ifwbqqsuppynhgp)**.

## Acknowledgments

This work has been supported financially by the National Science Centre, Poland (NCN) grant 2020/39/I/NZ2/02464 to P.A.Z. The computations were performed at the Poznan Supercomputing and Networking Centre (grant 312). W.D. and M.S.-L. were recipients of START 2023 Scholarship (Foundation for Polish Science). W.D. was a recipient of Best PhD Scholarship 2023 (AMU Foundation).

## Author contributions

W.D & P.A.Z. conceived the research idea. W.D., A.B., T.B., M.K., A.P., A.W., M.S.-L. performed experiments. All the authors participated in writing and correcting the manuscript.

## Conflict of interest

The authors declare no competing interest.

## Notes

### Competing Interest Statement

The authors have declared no competing interest.

