## Supplementary Figures for "Chromatin crosstalk between HDA19 and NuA4 sets thresholds for stress gene activation in *Arabidopsis*"

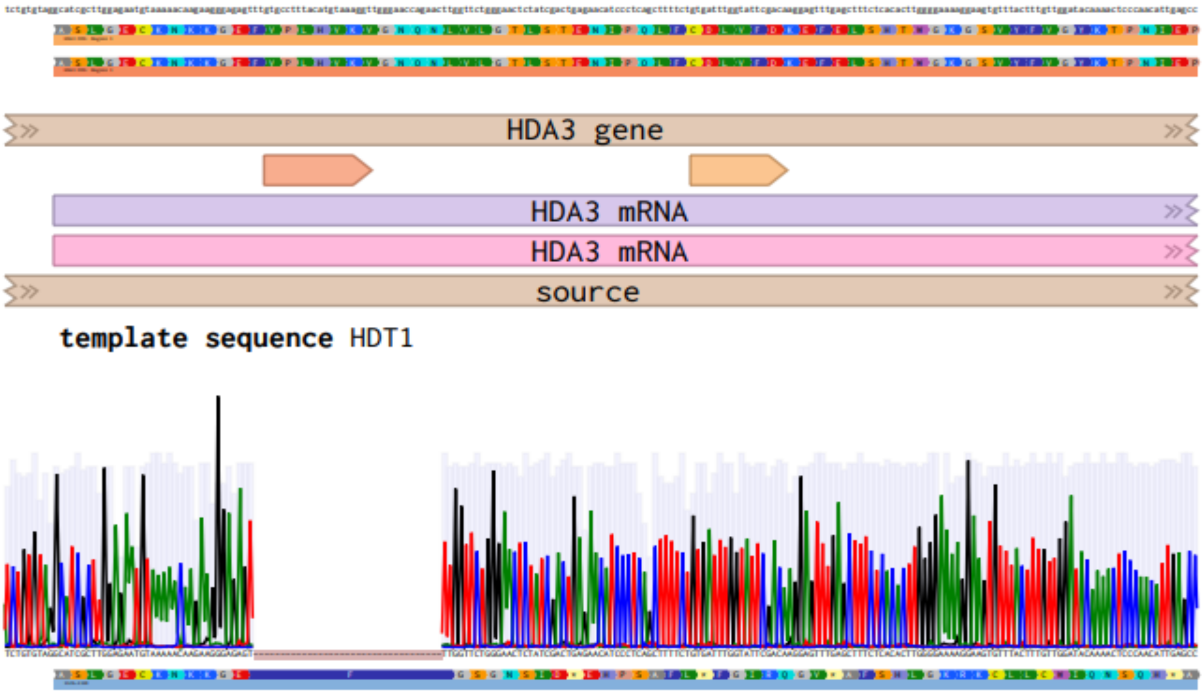


**Supplementary Figure 1.** Sanger sequencing result confirming out of frame deletion in *hdt1-3*. GuideRNA sequences are represented as 20 bp arrows above mRNA annotations. Stop codons are symbolized as asterisks on light-yellow background below the chromatograph.

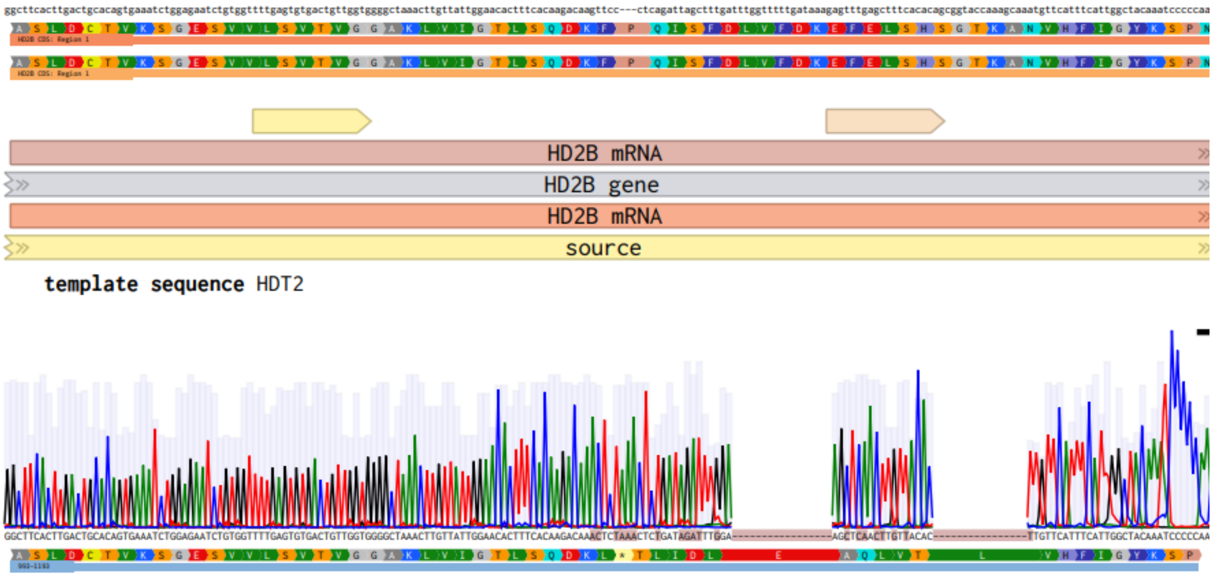


**Supplementary Figure 2.** Sanger sequencing result confirming out of frame deletion in *hdt2-3*. GuideRNA sequences are represented as 20 bp arrows above mRNA annotations. Stop codons are symbolized as asterisks on light-yellow background below the chromatograph.

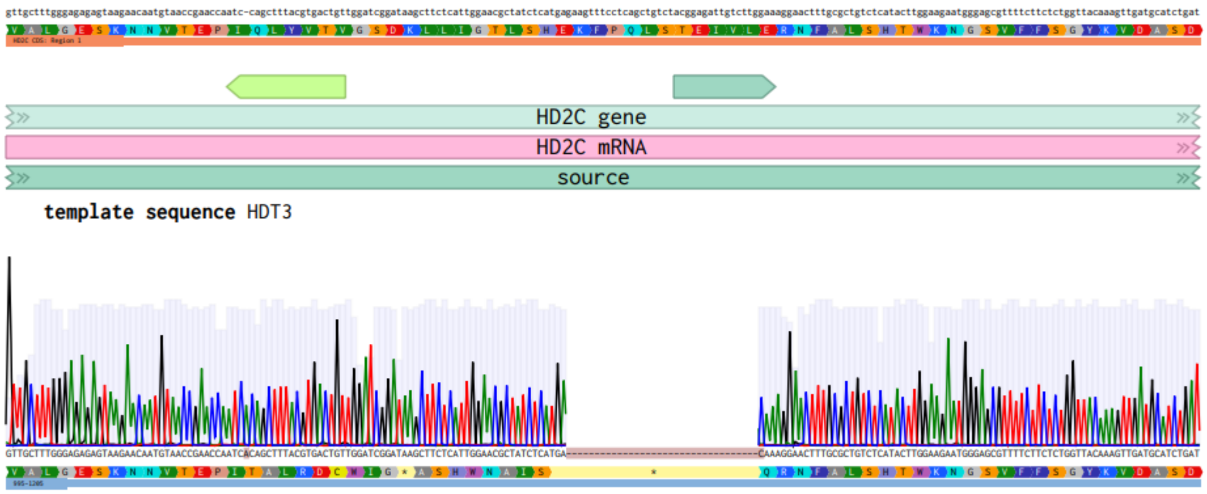


**Supplementary Figure 3.** Sanger sequencing result confirming out of frame deletion in *hdt3-3*. GuideRNA sequences are represented as 20 bp arrows above mRNA annotations. Stop codons are symbolized as asterisks on light-yellow background below the chromatograph.

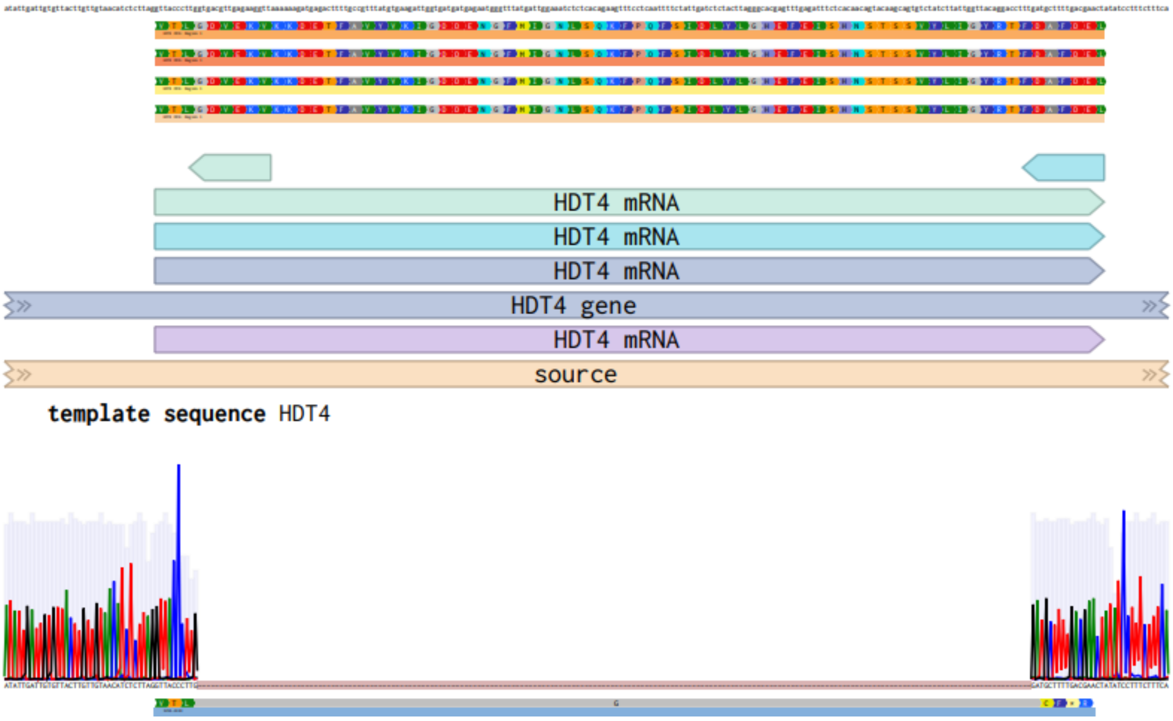


**Supplementary Figure 4.** Sanger sequencing result confirming out of frame deletion in *hdt4-2*. GuideRNA sequences are represented as 20 bp arrows above mRNA annotations. Stop codons are symbolized as asterisks on light-yellow background below the chromatograph.

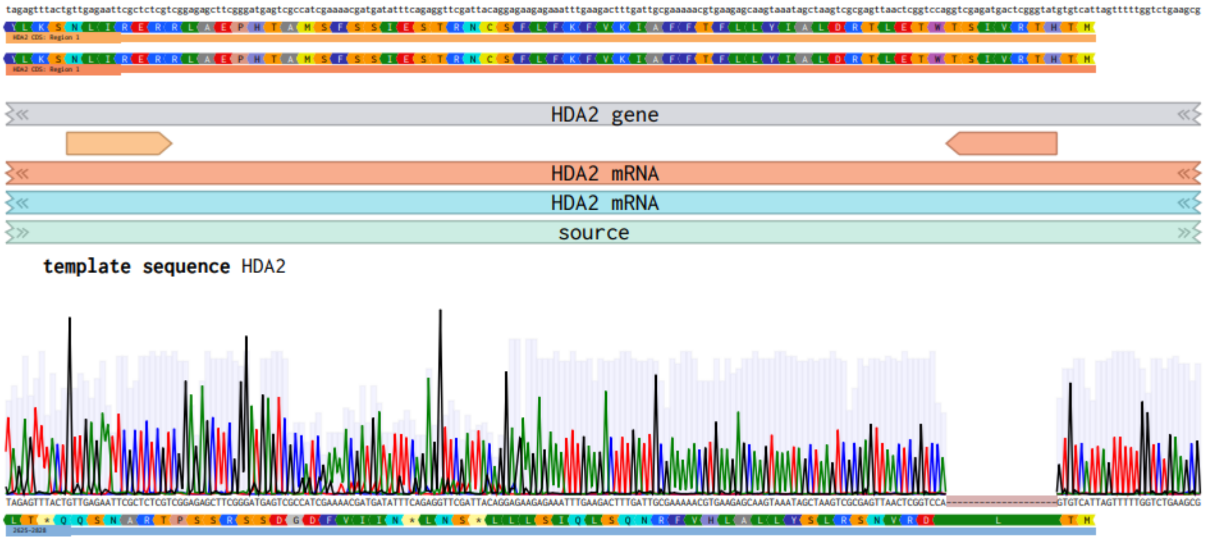


**Supplementary Figure 5.** Sanger sequencing result confirming out of frame deletion in *hda2-1*. GuideRNA sequences are represented as 20 bp arrows above mRNA annotations. Stop codons are symbolized as asterisks on light-yellow background below the chromatograph.

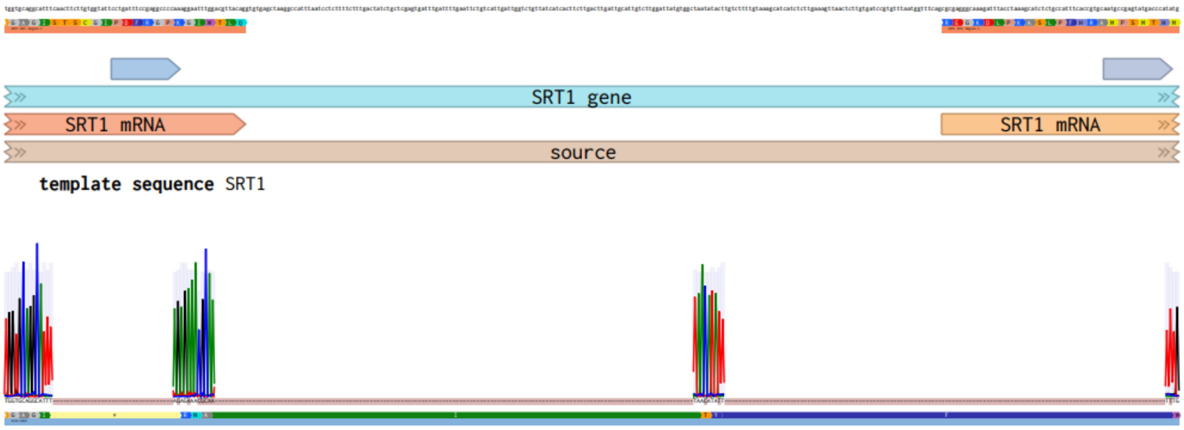


**Supplementary Figure 6.** Sanger sequencing result confirming out of frame deletion in *srt1-4*. GuideRNA sequences are represented as 20 bp arrows above mRNA annotations. Stop codons are symbolized as asterisks on light-yellow background below the chromatograph.

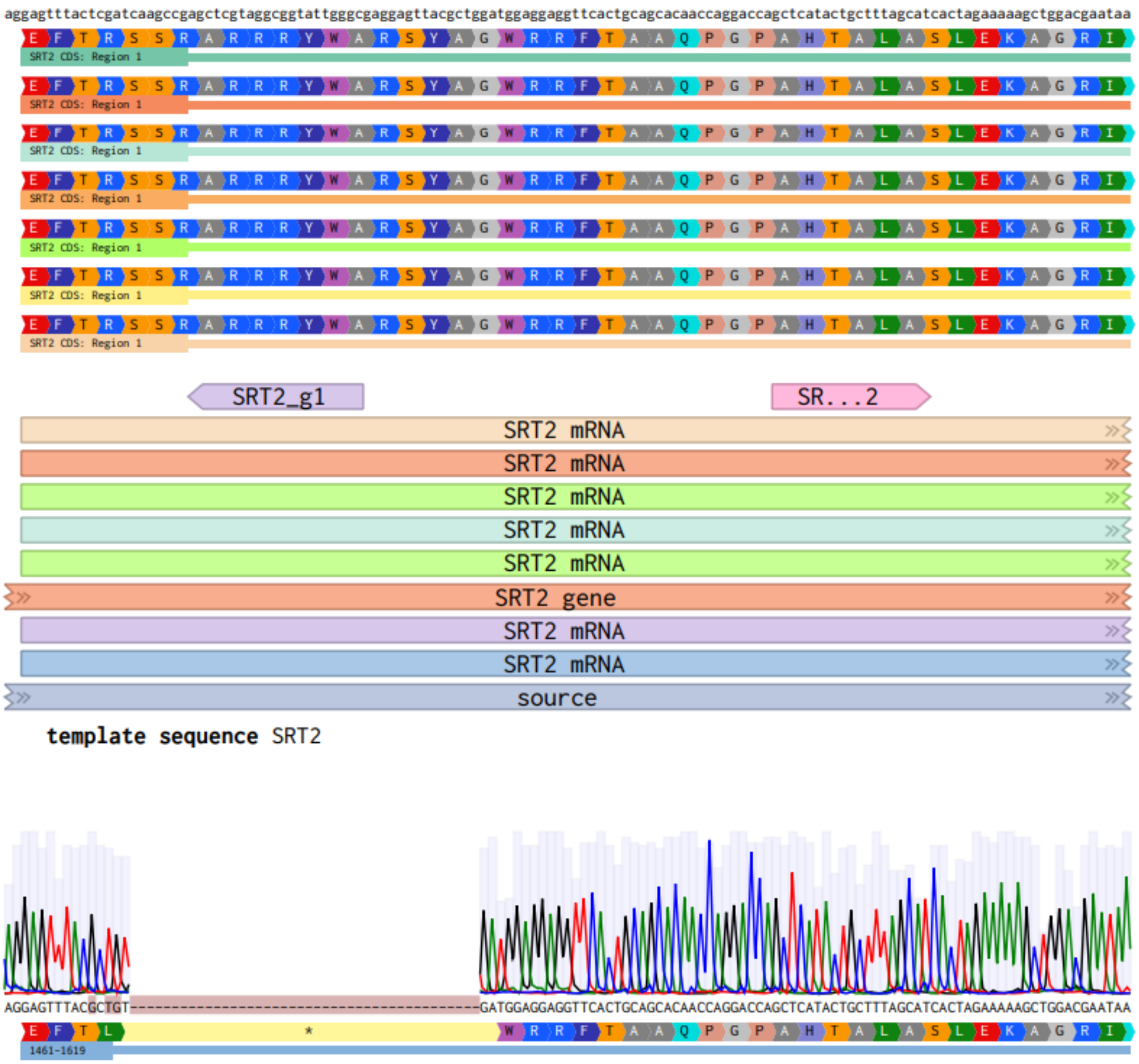


**Supplementary Figure 7.** Sanger sequencing result confirming out of frame deletion in *srt2-3*. GuideRNA sequences are represented as 20 bp arrows above mRNA annotations. Stop codons are symbolized as asterisks on light-yellow background below the chromatograph.

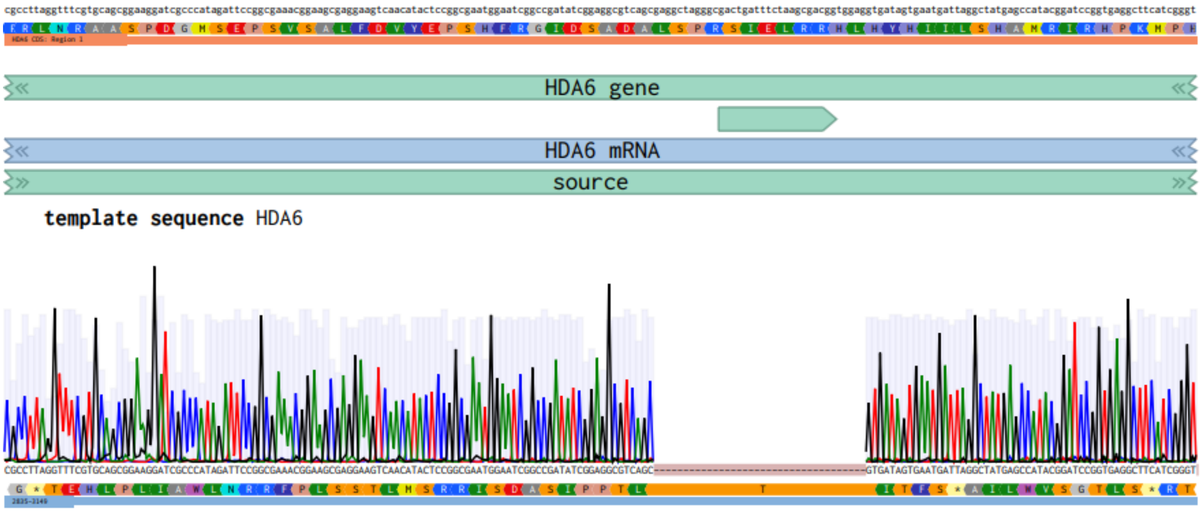


**Supplementary Figure 8.** Sanger sequencing result confirming out of frame deletion in *hda6-1*. GuideRNA sequence is represented as 20 bp arrow above mRNA annotations. Stop codons are symbolized as asterisks on light-yellow background below the chromatograph.

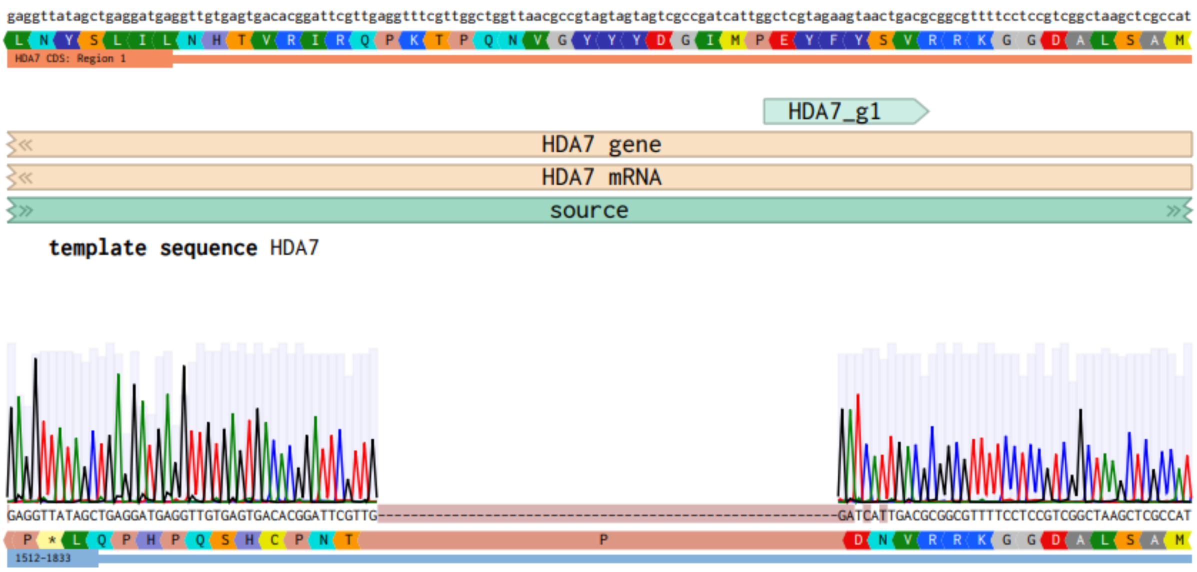


**Supplementary Figure 9.** Sanger sequencing result confirming out of frame deletion in *hda7-3*. GuideRNA sequences is represented as 20 bp arrow above mRNA annotations. Stop codons are symbolized as asterisks on light-yellow background below the chromatograph.

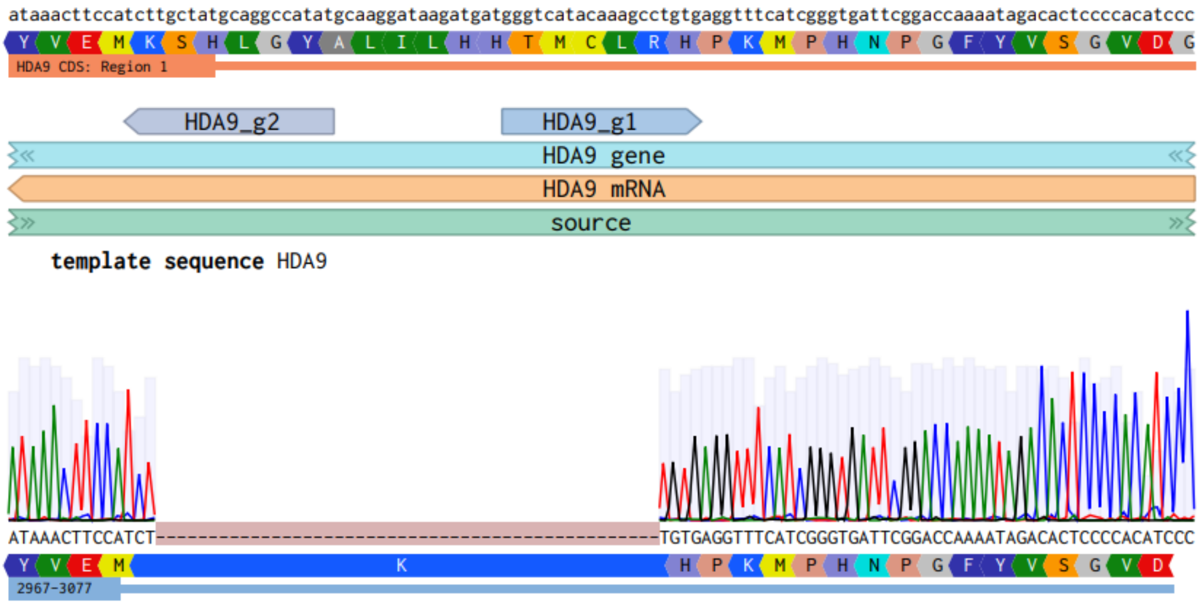


**Supplementary Figure 10.** Sanger sequencing result confirming in frame deletion in *hda9-3*. GuideRNA sequences are represented as 20 bp arrows above mRNA annotations.

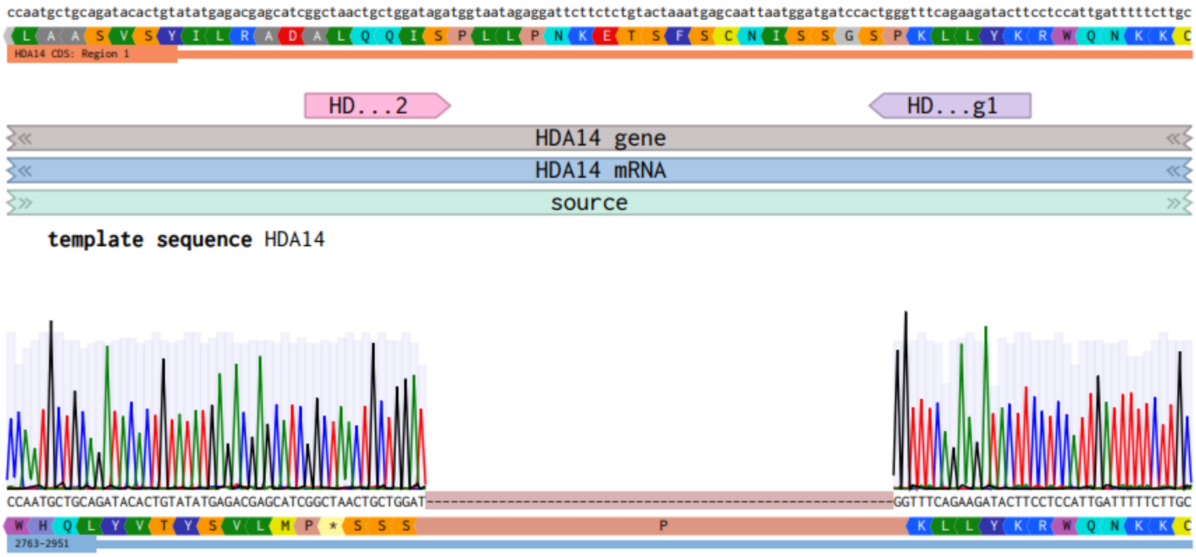


**Supplementary Figure 11.** Sanger sequencing result confirming out of frame deletion in *hda14-2*. GuideRNA sequences are represented as 20 bp arrows above mRNA annotations. Stop codons are symbolized as asterisks on light-yellow background below the chromatograph.

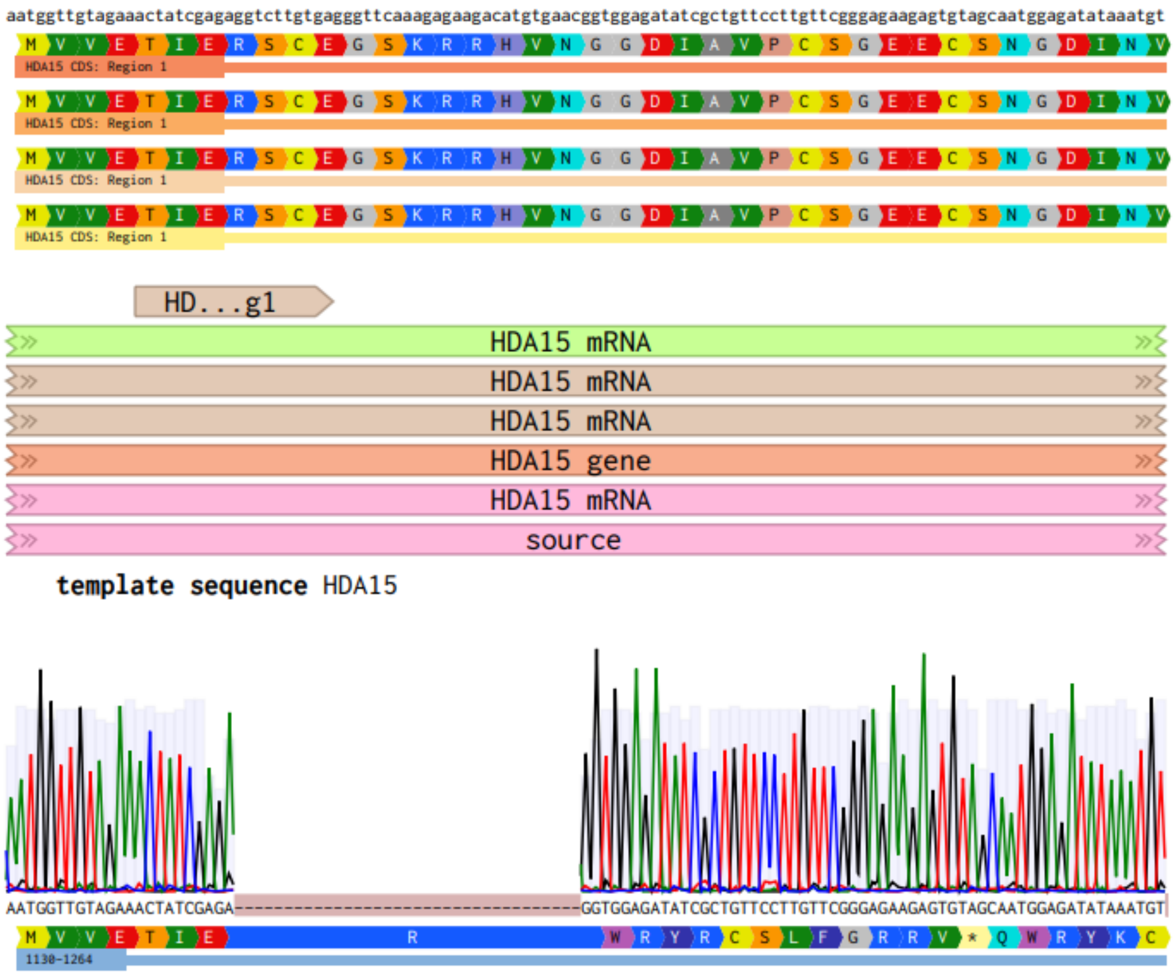


**Supplementary Figure 12.** Sanger sequencing result confirming out of frame deletion in *hda15-2*. GuideRNA sequences is represented as 20 bp arrow above mRNA annotations. Stop codons are symbolized as asterisks on light-yellow background below the chromatograph.

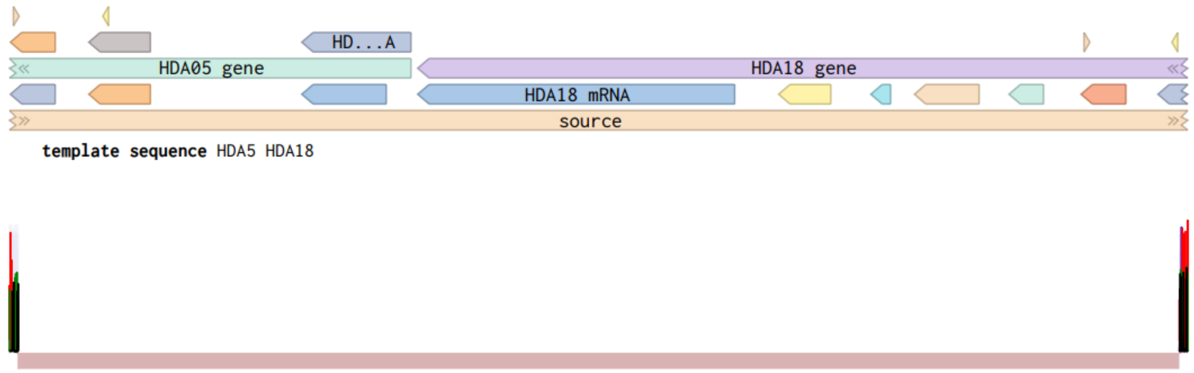


**Supplementary Figure 13.** Sanger sequencing result confirming large 3.8 kb deletion over tandemly duplicated *HDA5* and *HDA18* genes. GuideRNA sequences are represented as small yellow triangles above mRNA annotations.

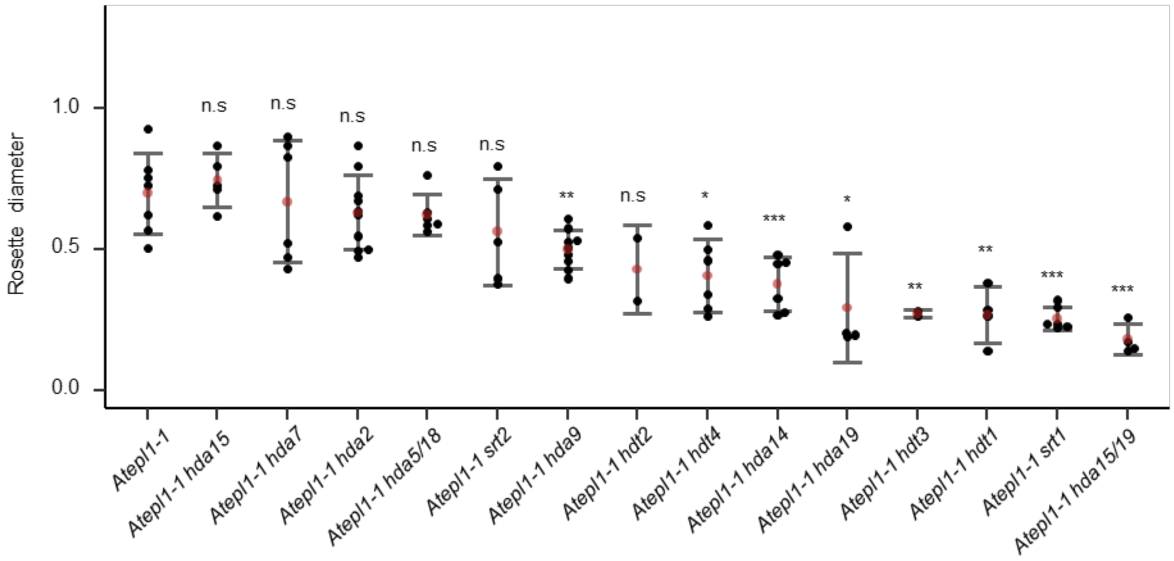


**Supplementary Figure 14.** Comparison of *Atepl1-1 hda* triple mutant rosettes, performed 28 DAG.

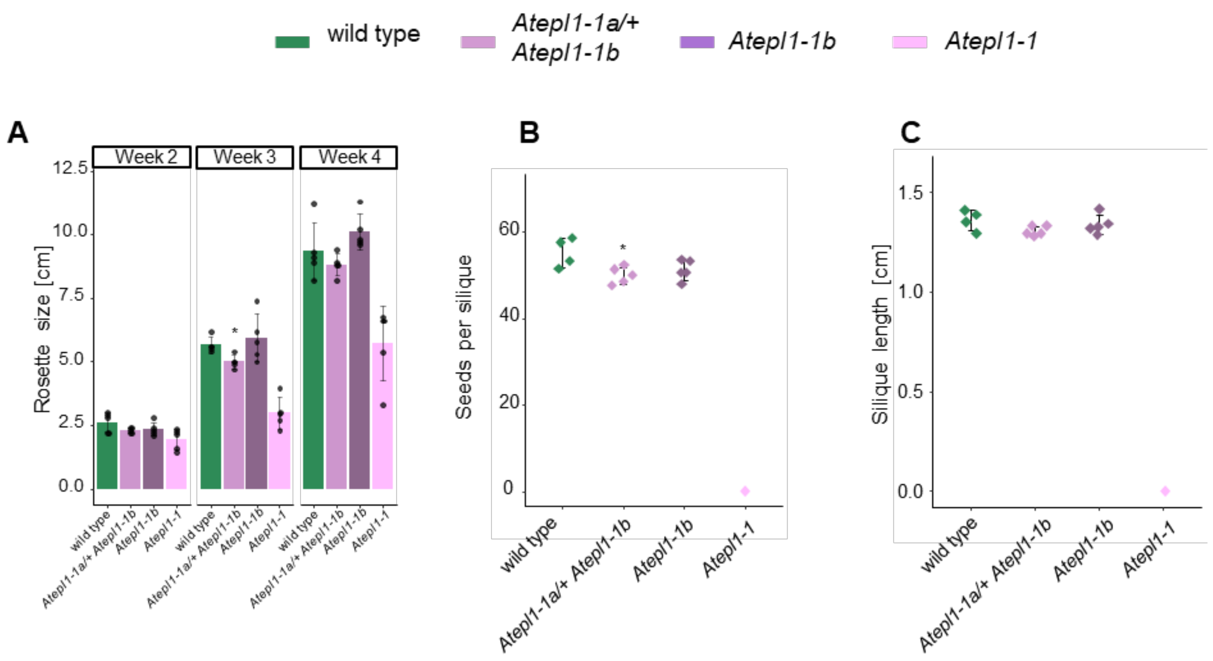


**Supplementary Figure 15**. Phenotype of *Atepl1-1* mutants. **A** Growth dynamics of wild type, *Atepl1-1a*/+ *Atepl1-1b*, *Atepl1-1b* and *Atepl1-1* (double mutant). **B** Fertility tests of wild type, *Atepl1-1a*/+ *Atepl1-1b*, *Atepl1-1b* and *Atepl1-1*. *Atepl1-1* mutant displays complete sterility. Data for *Atepl1-1* mutant incorporated from Bieluszewski et al., 2022.

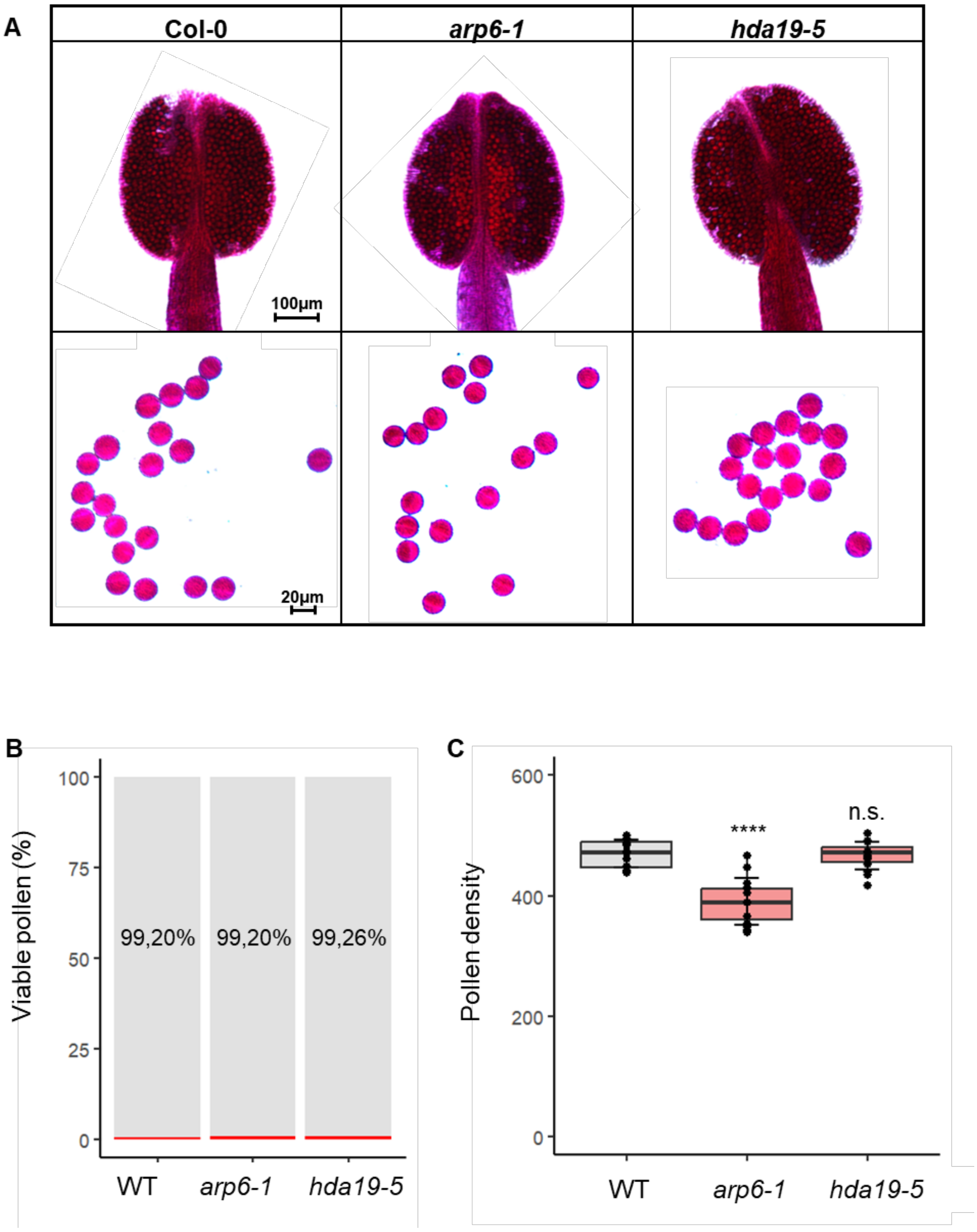


**Supplementary Figure 16**. Fertility of wild type (Col-0), *arp6-1* and *hda19-5* as visualized by Alexander staining. **A** Top panel shows representative picture of anthers. Scale bar, 100 µm. Bottom panel shows representative examples of pollen grains. Scale bar, 20 µm. **B** Pollen viability assessment of WT, *arp6-1* and *hda19-5* **C** Pollen density analysis for WT, *arp6-1* and *hda19-5*. Statistical significance was determined with two-sided Welch's t-test. Significance levels: $p < 0.0001$ ****, non-significant n.s.

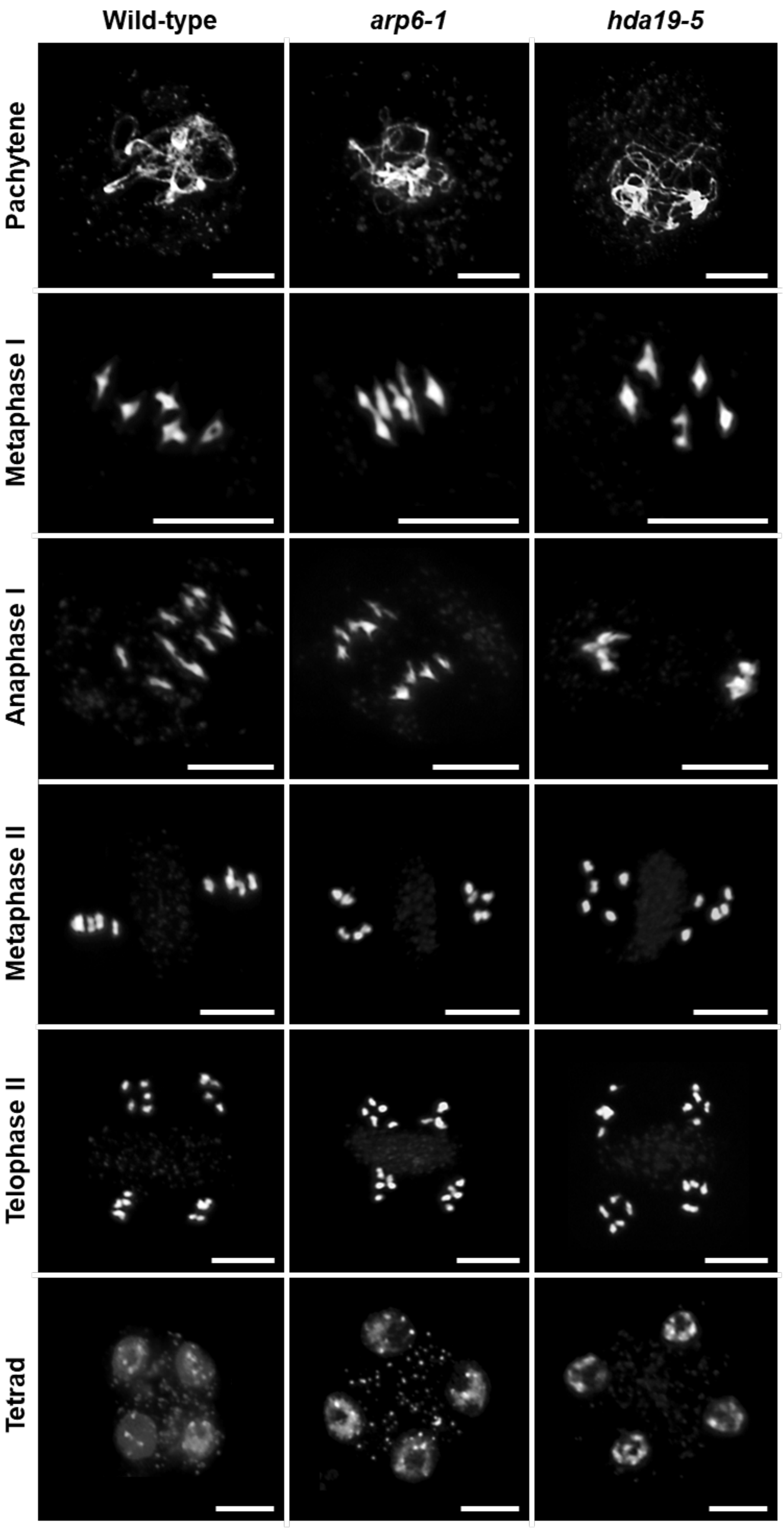


**Supplementary Figure 17.** Meiosis progression in *hda19-5* and *arp6-1* mutants.

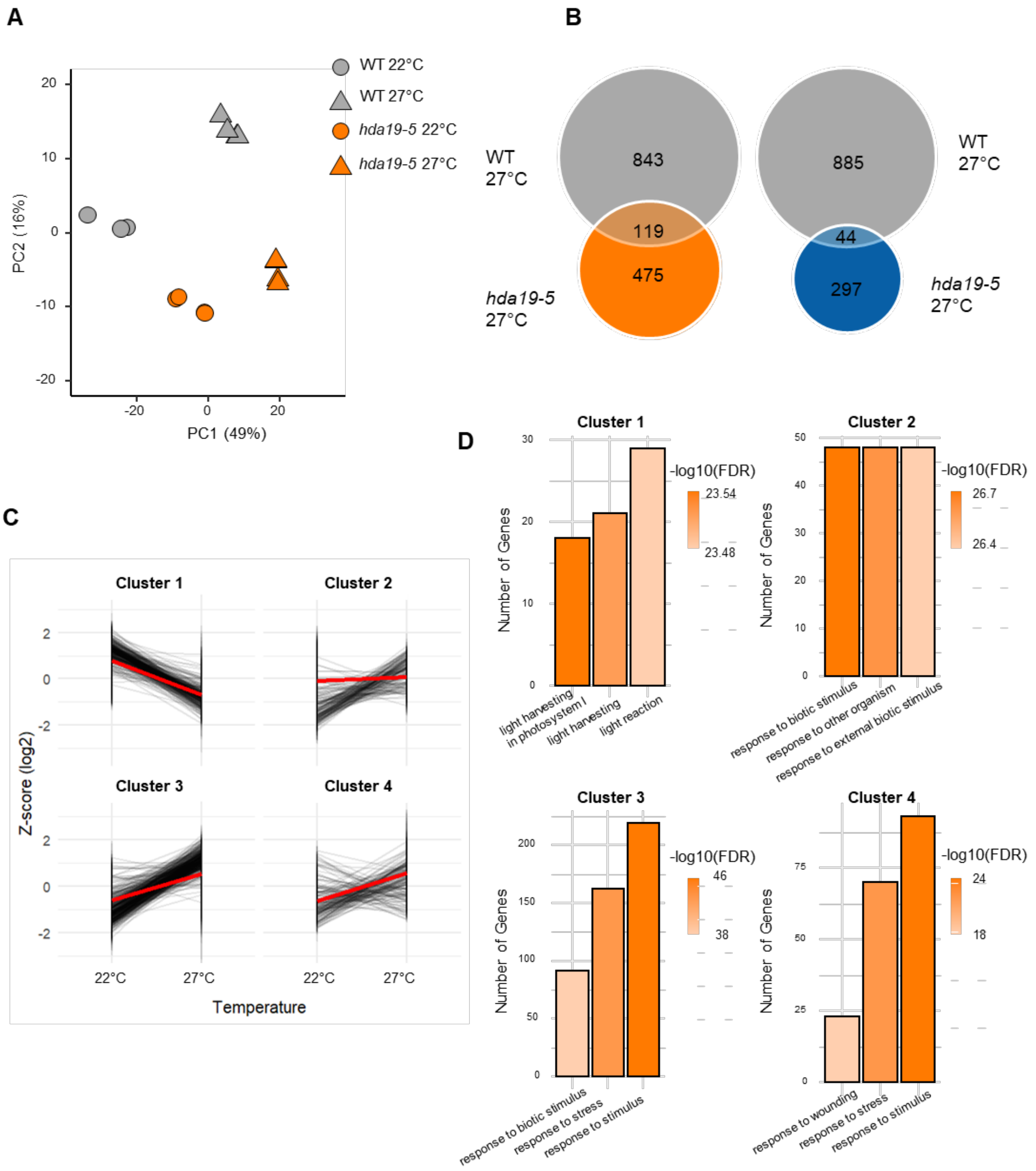


**Supplementary Figure 18.** Comparative analysis of 3'RNA-seq. **A** Principal component analysis of gene expression datasets from wild type (grey) and *hda19-5* (orange) mutants, grown at normal and elevated temperatures. **B** Venn diagrams showing overlaps between upregulated and downregulated genes between *hda19-5* and wild-type plants grown at 27°C. **C** Expression profiles of genes grouped in clusters after k-means clustering at 22°C and 27°C. **D** Gene ontology of top 3 terms, found within four clusters.

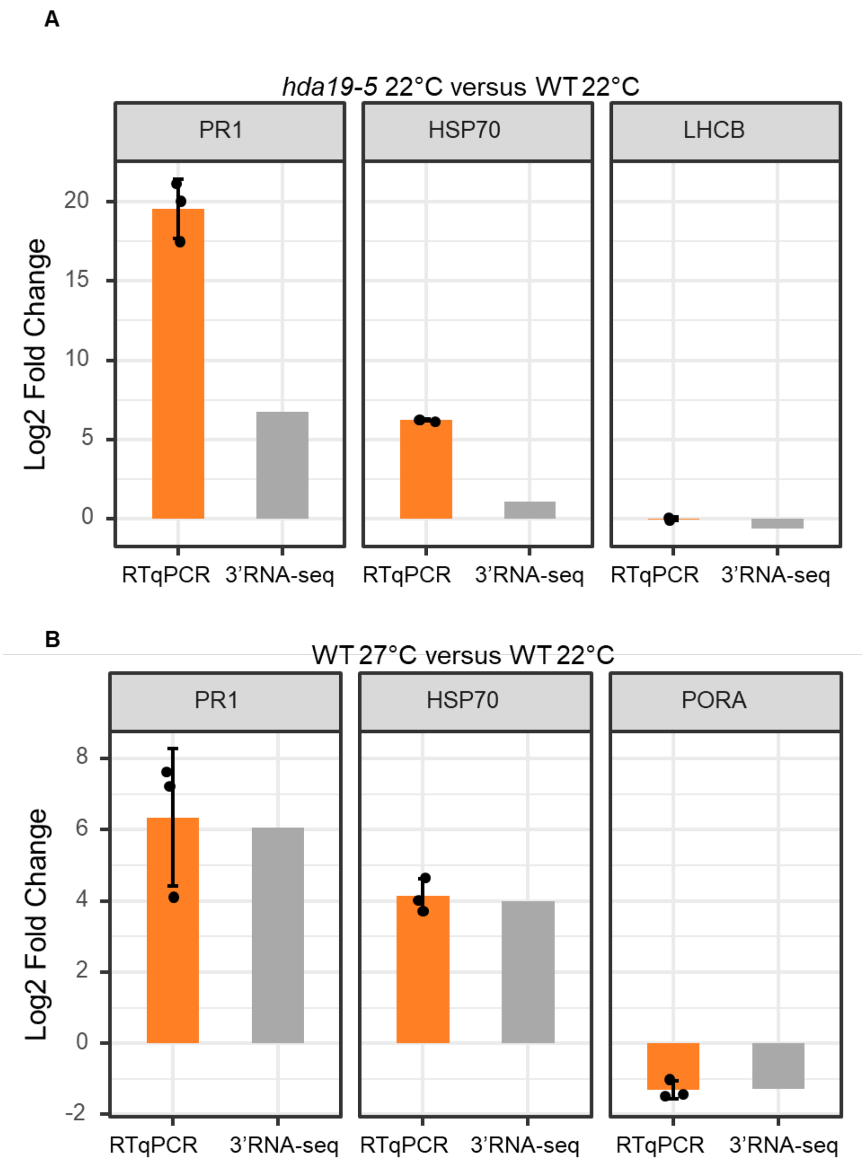


**Supplementary Figure 19.** Confirmation of 3′RNA-seq. **A** RTqPCR analysis of PR1, HSP70 and LHCB genes shows similar direction of expression change as previously reported in 3′RNA-seq in *hda19-5*. **B** Expression analysis of PR1, HSP70 and PORA genes in wild-type plants grown at 27°C with RTqPCR.

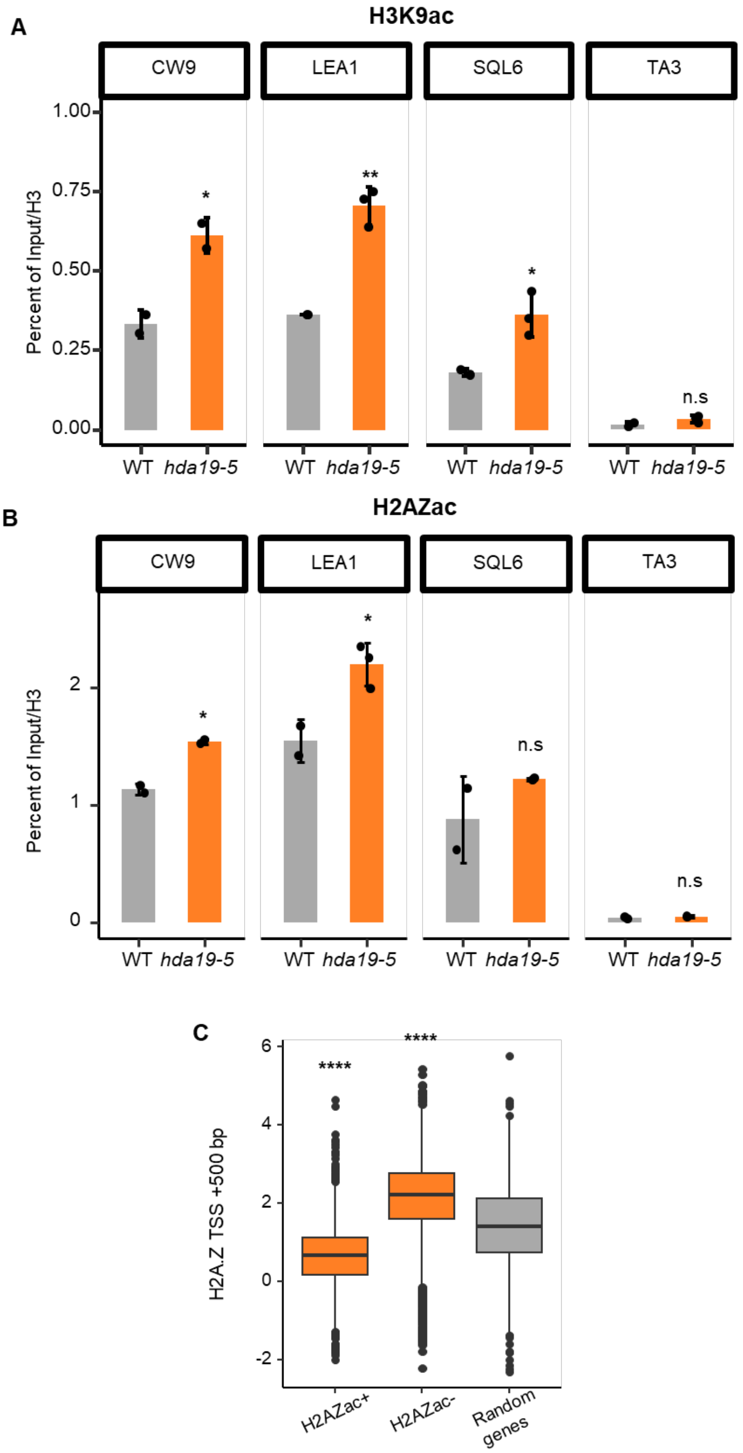


**Supplementary Figure 20.** Levels of H3K9ac and H2A.Zac measured by ChIP-qPCR. **A** ChIP-qPCR confirming gain of H3K9ac in 3 stress-related genes. Gypsy transposon (TA3; *AT5G15710*) was used as a negative control. **B** ChIP-qPCR confirming higher levels of H2A.Zac at 3 genes related to stress response. Gypsy transposon was used as a negative control. Statistical significance was determined with two-sided Welch's t-test. Significance levels: $p < 0.05$ *, $p < 0.01$ **, non-significant n.s. **C** Boxplot showing levels of total H2A.Z at +500 bp from TSS in the genes that gain H2A.Zac, loose H2A.Zac and 2500 random genes in the *hda19-5* mutant background. Statistical significance $p < 2.2e\text{-}16$ (****) was calculated between differentially enriched genes and random set of genes with use of Wilcoxon test.

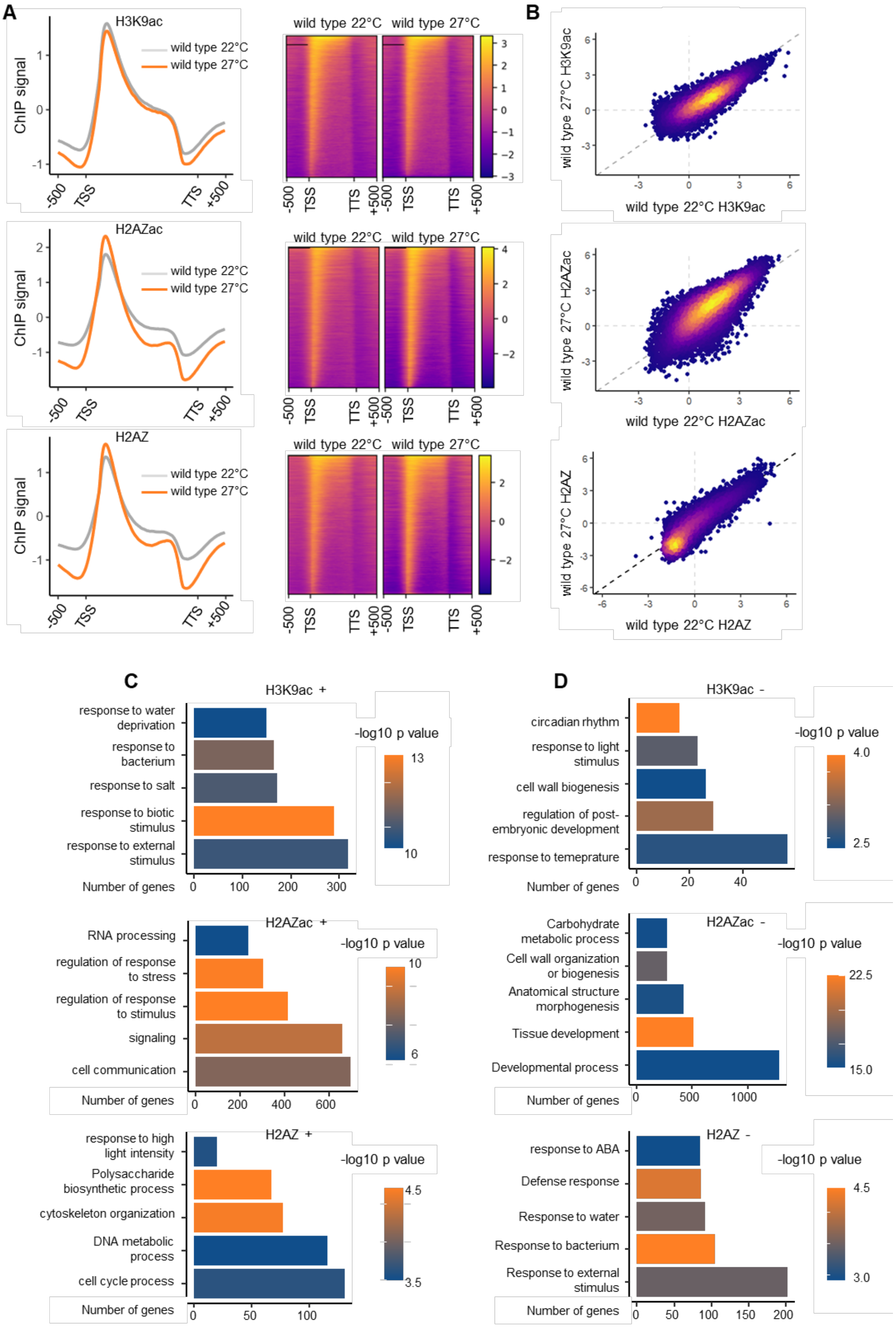


**Supplementary Figure 21.** Analysis of ChIP data for WT grown at 27°C. **A** Metagene H3K9ac signal plot for wild type grown at 22°C (grey) and wild type grown at 27°C (orange) expressed genes (n = 19,256; left). TSS marks transcription start site, TTS marks transcription termination site. Heatmaps displaying occupancy of H3K9ac at expressed genes in wild type grown at 22°C and wild type grown at 27°C. Amount of H3K9ac enrichment over input is colour encoded (right). Scatterplot for individual genes showing H3K9ac levels in wild type grown at 22°C and wild type grown at 27°C. Point density is colour encoded (right). The values on scatterplot (right) represent amounts of H2A.Z found in the gene body region. **C -D** Gene ontology analysis of genes showing differential enrichment in H3K9ac (top), H2A.Zac (middle) and H2A.Z (bottom). X-axis represents numbers of genes in each enriched category shown on y-axis. Color of the bars reflects statistical significance (-log10 p-value of FDR). 5 most unique and abundant categories were selected for visualization.

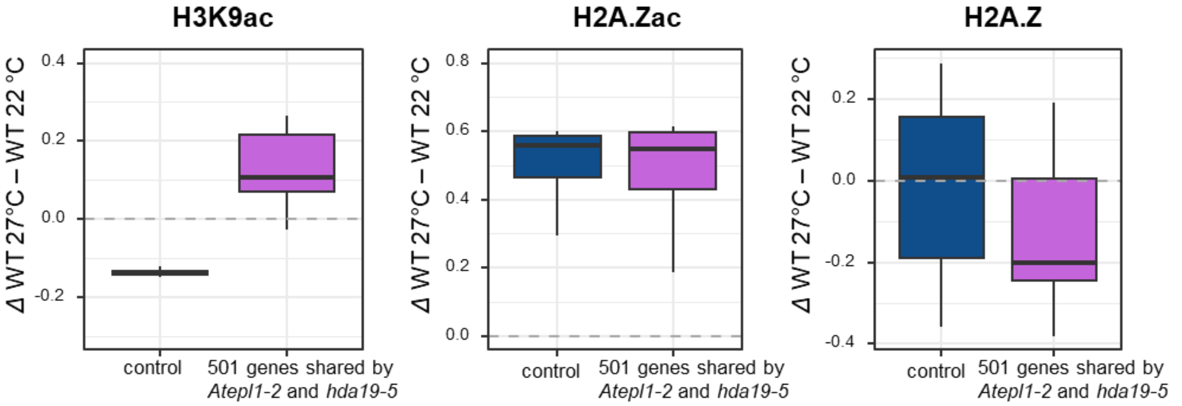


**Supplementary Figure 22**. HDA19-NuA4 controlled genes in wild-type grown at 27°C. Analysis of differences in H3K9ac, H2A.Zac and H2A.Z between wild type grown in physiological and elevated temperatures.
